# Testing microbial biomining from asteroidal material onboard the International Space Station

**DOI:** 10.1101/2024.01.13.575412

**Authors:** Rosa Santomartino, Giovanny Rodriguez Blanco, Alfred Gudgeon, Jason Hafner, Alessandro Stirpe, Martin Waterfall, Nicola Cayzer, Laetitia Pichevin, Gus Calder, Kyra R. Birkenfeld, Annemiek C. Waajen, Scott McLaughlin, Alessandro Mariani, Michele Balsamo, Gianluca Neri, Lorna J. Eades, Charles S. Cockell

## Abstract

Expanding human space exploration beyond Earth’s orbit necessitates efficient technologies for self-sustainable acquisition of local resources to overcome unviable resupply missions from Earth. Potential source of materials are asteroids, some of which contain valuable metals, such as platinum group elements.

The BioAsteroid experiment, performed onboard the International Space Station, tested the use of microorganisms (bacteria and fungi) to carry out mining of useful elements from asteroidal material (L-chondrite) under microgravity, in support of a long-term human presence in space. The fungus *Penicillium simplicissimum*, enhanced the mean release of palladium, platinum and other elements from the meteorite material in microgravity, compared to non-biological leaching. However, there was large variability in the results. For many elements, non-biological leaching under microgravity was enhanced compared to terrestrial gravity, while bioleaching was unaffected. Metabolomics results revealed clear patterns that highlight the influence of space conditions on the microbial metabolism, particularly for *P. simplicissimum*. We identified the presence of carboxylic acids, and molecules of potential biomining and pharmaceutical interest, enhanced in microgravity.

These results show a non-trivial effect of microgravity on bioleaching, highlighting the requirement of an optimal combination of microorganism(s), rock substrate, and conditions for successful biomining, both in space and Earth.

## 1. Introduction

To establish a long-term human presence in space, it will be necessary to extract resources from the local environment. This approach to space settlement, called *in situ* resource utilisation (ISRU), aims to reduce the mass and volume of resources that must be launched from Earth, and enable sustainable manufacturing of materials and products without the need for a mass and energy intensive constant resupply from Earth ^1–5^.

Asteroids, such as those in the asteroid belt and near-Earth asteroids ^2,6^, can be a potential source of elements and volatiles useful for future human space settlements, such as water, hydrogen, carbon compounds, metallic and non-metallic elements ^7–9^. Some asteroids are also known to contain high concentrations of precious metals including platinum-group elements (PGEs) ^8,10,11^. These, such as palladium and platinum, are used in high technology industries, and are of special interest because of their high melting points, corrosion resistance, and catalytic qualities ^12^. Although the economics of asteroid mining under the current technological development are yet to be determined ^13^, these elements command a high price on Earth, and have great potential use to support high technology manufacturing in space ^6,14^.

One possible way to extract elements from extraterrestrial materials is to use physico-chemical methods. However, in recent decades microorganisms have been recognised as an alternative and sustainable way to carry out, and efficiently catalyse, chemical transformations ^2,6,15^. For instance, it has been suggested that microorganisms could be used as the basis of a ‘bio-manufactory’ on Mars ^16,17^. Indeed, microorganisms are already employed in a wide range of manufacturing processes on Earth such as in the food, drugs, and chemical feedstock industries, and for further processing such as in plastics production. Similar versatilities could be applied in space ^18^, unlocking sustainable approaches for human space exploration ^4^.

One important process carried out by microorganisms is biomining, a technology where organisms are used to catalyse the breakdown of rocks and the release of useful elements, accelerating the acquisition of the required elements, exploiting mine waste tailings, and avoiding the use of environmentally damaging toxic compounds such as cyanides ^19^. Biomining is a widely adopted process on Earth, for instance to extract copper and gold ^20–23^. Well studied acidophilic iron and sulphur-oxidisers are often used to bioleach sulfidic ores^21,24,25^, but heterotrophic microorganisms, including bacteria and fungi^26–28^, are effective in bioleaching in environments with circumneutral pH. Their bioleaching capacity is enabled by the release of protons or organic acids (e.g.citric, oxalic,glucuronic acids, etc), thereby decreasing the pH in the system, or by the release of complexing compounds ^29^.

Biomining is a promising technology to extract useful elements and compounds from local materials in space ^2,6,30^. For example, the bacterium *Sphingomonas desiccabilis* was recently used to catalyse the extraction of rare earth elements and vanadium from basalt rock under microgravity and Martian gravity on the International Space Station (ISS) ^31,32^. Previously, bacterial leaching of copper has been demonstrated in microgravity ^33^, as well as bioleaching of elements from lunar and Martian simulants on Earth ^34,35^. Aside from mining, microbial interactions with regolith (loose material covering solid rocks on a planetary body surface) will be the first stage of breaking down rocks to make soils or to release nutrients in life support systems that employ regolith as a feedstock ^4^. Extraterrestrial materials, such as carbonaceous chondrite, have been shown to support the growth of microorganisms^36–38^.

In this study (BioAsteroid), we showed that heterotrophic microorganisms (the bacterium *S. desiccabilis*^31,32,39,40^ and the fungus *Penicillium simplicissimum*^23,41–43^) can be used to catalyse the release of technologically and economically important elements from L-chondrite material, a common type of meteorite^8^, under microgravity conditions onboard the International Space Station (ISS; Figure 1). Microbial consortia are often beneficial in terrestrial biomining^22,44^. For this reason, we augmented single-species samples with a consortium formed by the two microorganisms. We focused on bioleaching of three PGEs and other 41 elements of industrial interest, highlighting the effect of the organism compared to abiotic leaching, and the effect of microgravity compared to Earth conditions. A thorough metabolomic analysis allowed us to study the metabolic responses to microgravity during microbe-mineral interactions. These experiments demonstrate proof of principle for the use of microorganisms to transform asteroidal material for future human exploration and settlement of space in a self-sustainable fashion.

**Figure 1.**
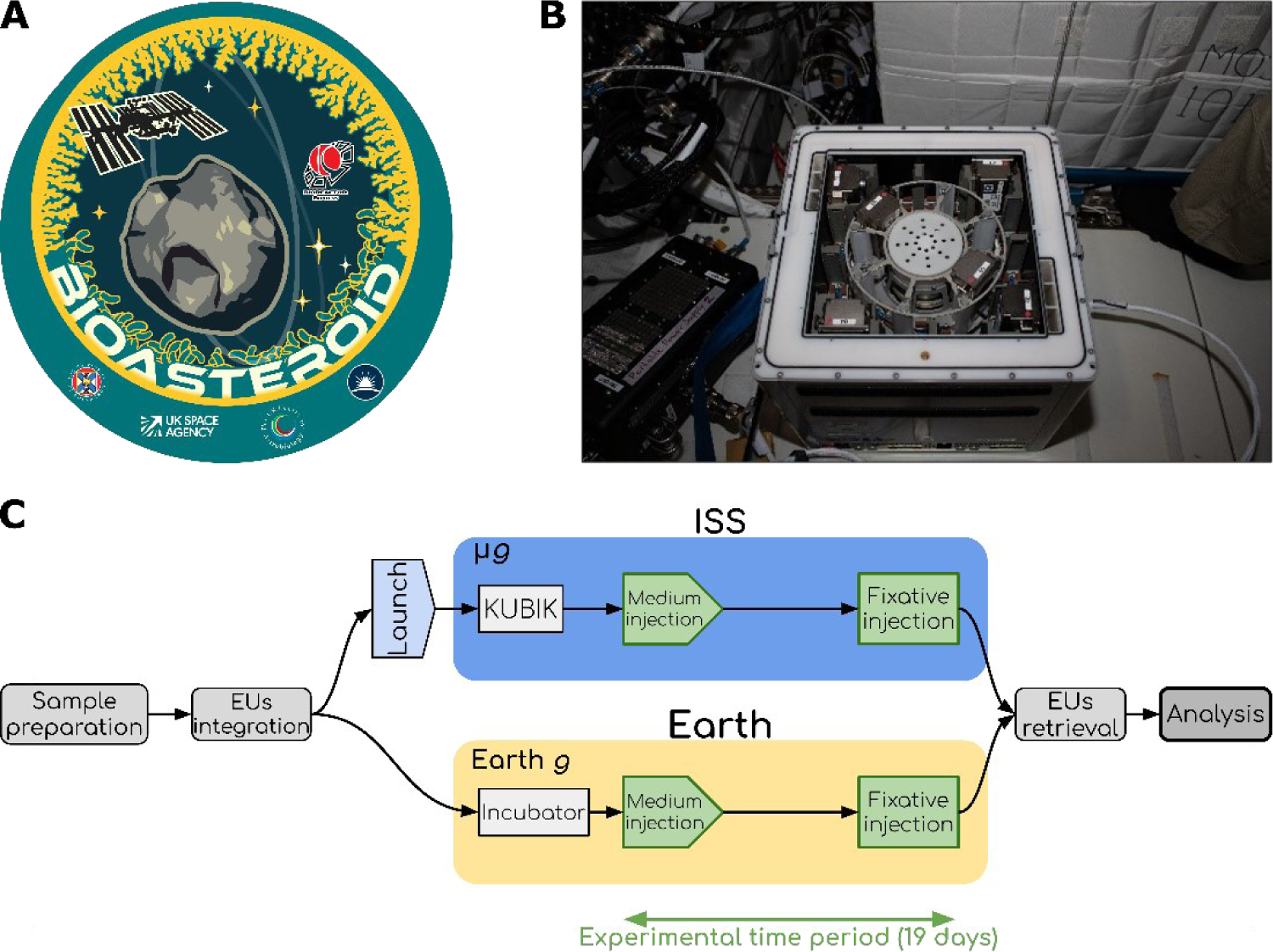
The BioAsteroid experiment. A) BioAsteroid logo, created by Sean McMahon (University of Edinburgh); B) The six hardware units inserted into the KUBIK onboard the ISS (credits ESA/NASA); C) Flow diagram of the experiment. After preparation, samples were integrated into the experimental units (EUs) together with the medium and the fixative. The EUs were either launched to the ISS (blue oval), where they were installed in KUBIK incubators and subjected to microgravity (µ*g*) or kept for incubation on Earth for the terrestrial gravity control (Earth *g*, yellow oval). Steps in green were part of the experimental time period (19 days). Storage passages were omitted for brevity.

## 2. Results

### 2.1 Meteorite characterisation

In order to identify the minerals and elements available for leaching, we thoroughly characterised the L-chondrite used in this experiment. Phase composition analysis was performed by X-ray Diffraction Spectroscopy (XRD), Raman spectroscopy, and backscatter electron microscopy (BSE) with energy dispersive spectrometry (EDS) elemental mapping. Inductively coupled plasma mass spectrometry (ICP-MS), and inductively coupled plasma optical emission spectroscopy (ICP-OES) were used to determine elemental availability to the microbial cells (Figure 2, Tables 1-2, S1).

**Figure 2.**
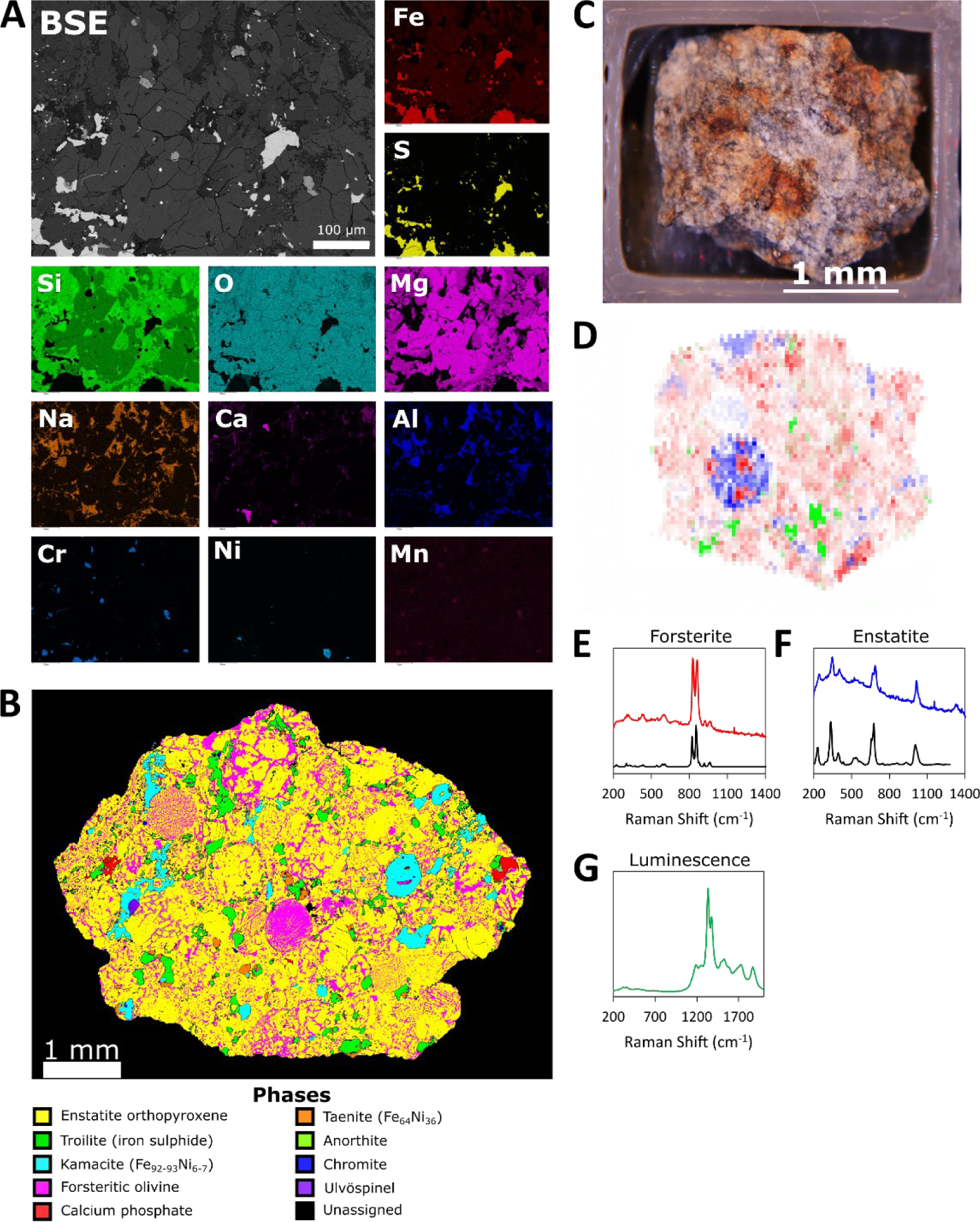
Characterization of the L-chondrite meteorite. A) Backscatter electron image (BSE) and single elemental mapping for some of the major elements present in a pristine representative fragment of the L-chondrite used in this experiment (scale bar: 100 µm); B) Phase analysis of a similar pristine fragment of the L-chondrite; C) photographic image of a pristine fragment analysed by Raman spectroscopy; D) composite Raman map with forsterite in red, enstatite in blue, and luminescence in green. Typical spectra are displayed for E) forsterite, F) enstatite, and G) luminescence signal. Reference spectra (black) are displayed for the minerals.

**Table 1.**
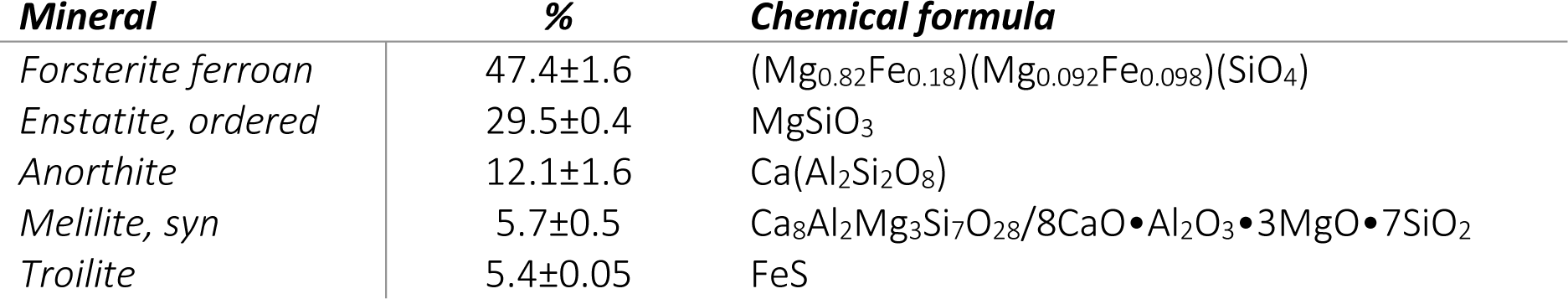
XRD analysis of pristine meteorite fragments, indicating percentage mineral composition (mean±st. error); n=3.

| **Mineral** | **%** | **Chemical formula** |
| --- | --- | --- |
| *Forsterite ferroan* | 47.4±1.6 | (Mg<sub>0.82</sub>Fe<sub>0.18</sub>)(Mg<sub>0.092</sub>Fe<sub>0.098</sub>)(SiO<sub>4</sub>) |
| *Enstatite, ordered* | 29.5±0.4 | MgSiO<sub>3</sub> |
| *Anorthite* | 12.1±1.6 | Ca(Al<sub>2</sub>Si<sub>2</sub>O<sub>8</sub>) |
| *Melilite, syn* | 5.7±0.5 | Ca<sub>8</sub>Al<sub>2</sub>Mg<sub>3</sub>Si<sub>7</sub>O<sub>28</sub>/8CaO•Al<sub>2</sub>O<sub>3</sub>•3MgO•7SiO<sub>2</sub> |
| *Troilite* | 5.4±0.05 | FeS |

**Table 2.** ICP-MS and ICP-OES analysis of pristine meteorite fragments, indicating the concentration of various elements (mean±st. error); n=3. Elements are listed in order of abundance in the meteorite. Elements in pale yellow are PGEs.

| **ICP-MS** |  | **ICP-OES** |  |
| --- | --- | --- | --- |
| **Element** | **µg/g** | **Element** | **mg/g** |
| **Mo** | 6.032±2.510 | **Mg** | 159.399±18.262 |
| **Ce** | 1.122±0.032 | **Fe** | 84.502±8.279 |
| **Nd** | 0.809±0.027 | **S** | 28.555±2.956 |
| **Pb** | 0.551±0.079 | **Mn** | 25.440±2.561 |
| **La** | 0.437±0.012 | **Ca** | 13.897±1.389 |
| **Dy** | 0.248±0.009 | **Na** | 8.001±0.797 |
| **Gd** | 0.196±0.006 | **Al** | 7.786±2.838 |
| **Ir** | 0.186±0.017 | **Ni** | 5.746±0.312 |
| **Yb** | 0.168±0.005 | **K** | 1.457±0.091 |
| **Pr** | 0.164±0.005 | **P** | 1.113±0.130 |
| **Er** | 0.161±0.005 | **Cr** | 1.015±0.059 |
| **Sm** | 0.145±0.005 | **Ti** | 0.677±0.101 |
| **Hf** | 0.112±0.005 | **Co** | 0.152±0.023 |
| **Pt** | 0.091±0.009 | **Zn** | 0.079±0.006 |
| **Ho** | 0.054±0.002 | **Cu** | 0.063±0.004 |
| **Eu** | 0.053±0.002 | **Sr** | 0.018±0.002 |
| **Rh** | 0.051±0.005 | **Cd** | 0.015±0.001 |
| **Th** | 0.049±0.001 | **Zr** | 0.013±0.001 |
| **Tb** | 0.037±0.001 | **Ba** | 0.008±0.001 |
| **Ta** | 0.032±0.001 | **Pb** | 0.007±0.000 |
| **Lu** | 0.025±0.001 | **Ag** | 0.000±0.000 |
| **Tm** | 0.025±0.001 |  |  |
| **U** | 0.019±0.001 |  |  |
| **Os** | 0.017±0.004 |  |  |
| **Hg** | 0.004±0.001 |  |  |
| **Tl** | 0.003±0.000 |  |  |
| **Pd** | 0.002±0.000 |  |  |
| **Ru** | 0.001±0.000 |  |  |

The XRD results of major (>5%) crystalline mineral types are shown in Table 1. The L-chondrite material used in the experiments shows a bulk composition typical for this meteoritic material ^8^. The mineralogy is dominated by olivine (the magnesium-rich forsterite end member) and secondarily by pyroxene (magnesium-rich end member enstatite). Minor contributions are found from feldspar (anorthite), melilite (sometimes associated with calcium–aluminium-rich inclusions in chondritic meteorites) and iron sulphides (toilite). XRD does not reveal solid metal inclusions, but microscopy showed the presence of iron-nickel inclusions (Figure 2A-B). Raman spectra were recorded at an array of points across the surface to identify minerals and map their distribution. Forsterite and enstatite were detected, as expected, and were heterogeneously distributed through the material (Figure 2C-G), showing that the microbial population is exposed to a heterogenous matrix of material. Sharp luminescence peaks were observed between 800 nm and 900 nm, and can be attributed to rare earth metal dopants in the mineral lattice (2G) ^45,46^.

ICP-MS and ICP-OES (Table 2) showed that the most abundant element is magnesium (159.399±18.262 mg/g), followed respectively by iron (84.502±8.279 mg/g), sulphur (28.555±2.956 mg/g), manganese (25.440±2.561 mg/g), calcium (13.897±1.389 mg/g), sodium (8.001±0.797 mg/g), and aluminium (7.786±2.838 mg/g). The elemental composition is consistent with elements and mineral phases identified by BSE and XRD (Table 1, Figure 2B).

### 2.2 Microbial final cell concentration and pH

Final cell concentration was measured immediately after sample recovery, as an indication of microbial growth, by analysing the liquid fraction of each sample by spectrometric analysis (optical density, *λ* = 600 nm), flow cytometry and colony forming unit (CFU) assay.

Optical density data at λ=600 nm (Figure 3A, S2) of the liquid fraction were obtained after spaceflight. All samples showed higher final cell concentration on Earth compared to space (ISS). Using a wavelength of 530 nm (S2), which is sometimes used for filamentous fungi ^47^, we observed a similar trend.

**Figure 3.**
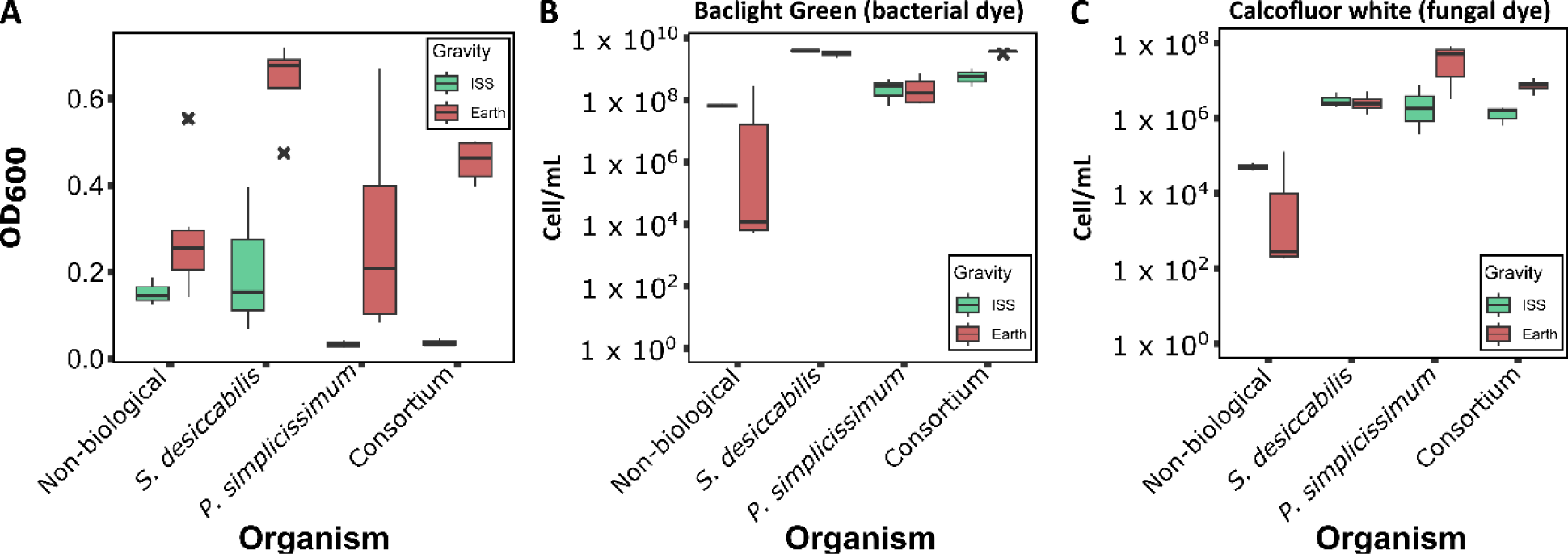
Final cell concentration in the liquid fraction of the BioAsteroid experiment. Optical density (*λ* = 600nm, A) and flow cytometry analysis measured (B-C) from the liquid fraction of the ISS and Earth samples. For flow cytometry, values represent the average events counted in 100 µL of sample stained with B) Baclight Green (specific for bacterial cells) and C) Calcofluor White (specific for the fungal cells). Black X represent outliers; n=3 for ISS samples, 4≤n≤6 for Earth samples.

Optical density at different wavelengths could potentially be sensitive to metabolic changes, therefore analysis of post-spaceflight final cell concentration was also obtained by flow cytometry (Figure 3B-C, S2). The flow cytometry results showed no gravity-driven difference between events measured with the bacterial dye for *S. desiccabilis* samples [(3.97±0.06)x10^9^ cell/mL on ISS, (3.27±0.32)x10^9^ cell/mL on Earth, respectively], and a slight increase (<10-fold) in Consortium samples (samples containing both the bacterium and the fungus) in Earth gravity compared to microgravity [(6.47±1.98)x10^8^ cell/mL on ISS against (3.64±0.14)x10^9^ cell/mL on Earth, respectively] (Figure 3B, S2). Differences in the fungi-associated events were present, when comparing *P. simplicissimum* in different gravities, with Earth samples showing concentrations one order of magnitude higher than the ISS [(3.50±1.99)x10^6^ cell/mL on ISS against (6.37±2.11)x10^7^ cell/mL on Earth, respectively]. Similarly, Consortium samples showed a slight increase (<10-fold) in final cell concentration on Earth samples compared to ISS [(1.43±0.34)x10^6^ cell/mL on ISS against (7.69±1.33)x10^6^ cell/mL on Earth, respectively]. The presence of positive events in the sterile diluent and non-biological controls (S2), potentially caused by small non-biological particulate matter, were in the worst case one order of magnitude smaller than the biological samples (<10%) and are thus considered negligible.

Additionally, CFU assay was performed to identify potential contaminations not identifiable by the previous two methods (S3). CFU analysis of *S. desiccabilis* showed reduced colony numbers in ISS compared to Earth samples, in both bacterial-only [(7.17 ±2.88)x10^6^ CFU/mL on ISS, (1.75±1.24)x10^11^ CFU/mL on Earth], and Consortium samples [(4.39±2.38)x10^6^ CFU/mL on ISS against (2.38±2.05)x10^10^ CFU/mL on Earth]. This is in contrast with the flow cytometry results for *S. desiccabilis*, but not for the other samples. In contrast, fungal CFU analysis of *P. simplicissimum* and Consortium showed concentrations two orders of magnitude higher on the ISS compared to Earth [*P. simplicissimum*: (1.37±1.01)x10^4^ CFU/mL on ISS, (4.58±3.08)x10^2^ CFU/mL on Earth; Consortium: (8.89 ±6.59)x10^3^ CFU/mL on ISS, (8.33±4.17)x10^1^ CFU/mL on Earth]. This is in contrast with both the OD_600_ and the flow cytometry measurements (Figure 3, S2). The discrepancies observed might be explained by the fact that CFU assay measures actively dividing cells, potentially reflecting different survival rates on Earth and ISS populations.

We observed contaminations in some of our CFU assays from endogenous (*S. desiccabilis*) and exogenous (Contaminant 1 and 2, S3) species. 16S rDNA sequencing of the contaminated ISS samples could not identify any species, suggesting the contamination was not present in the original samples, as corroborated by flow cytometry (Figure 3B). 16S rDNA sequencing of the contaminated Earth samples identified a species of the *Sphingomonas* genus for one of the two bacterial samples affected, and for two of the three Consortium samples affected. Identification failed for the remaining affected samples (see materials and methods for details). Taken together, this suggests original bacterial and Consortium Earth samples were not contaminated, or that any contamination was dominated by *S. desiccabilis*, in accordance with the CFU assay (S3). For the fungal Earth samples, alignment for three of the four affected sample sequences suggested a contamination from the order Bacillales, probably from the genera *Bacillus* and/or *Planococcus*. Identification failed for the fourth sample. Sequences of the isolated species (Contaminant 1 and 2) aligned with the genera *Bacillus* and *Paenibacillus*, suggesting these as the most probable genera for the contaminants. The presence of the contaminations was considered when interpreting the results. Spare samples prepared for the Earth control experiment did not show Contaminant 1 or 2 colonies formation (data not shown), suggesting the contamination did not occur during sample preparation but probably during sample post-flight processing or hardware integration.

The pH is one of the potential influencing factors in leaching. We measured the pH of the liquid fractions after spaceflight (S4), reporting values between 7.21 and 7.43 among all samples. It must be noted that the values were likely affected by the presence of the fixative (RNAlater, pH=4.87±0.01), which was necessary to halt the experiment at its end, and prevented us from deriving conclusions on the changing pH during the experiment.

### 2.3 Microbe-mineral interaction

To investigate if microgravity had an effect on the interaction of the microorganisms with the L-chondrite, and reveal if local mineral composition could influence it, microbe-mineral interaction was investigated using Scanning Electron Microscopy (SEM) secondary electron imaging (Figure 4) and EDS (S5-7) analysis to qualitatively assess the potential microbial preference for a particular mineral.

**Figure 4.**
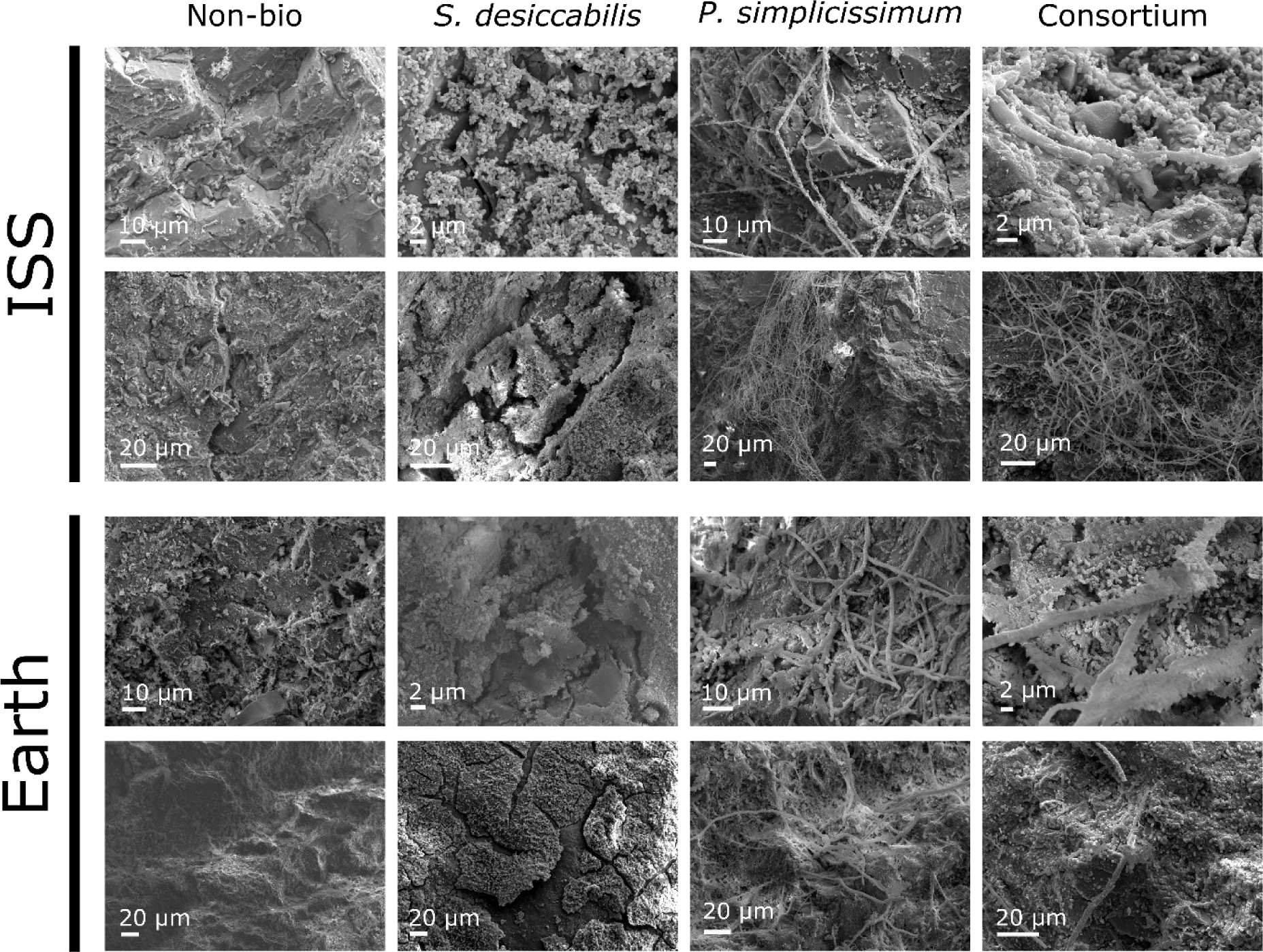
Scanning electron microscopy (SEM) images of the L-chondrite fragments. Secondary electron SEM images are shown here and are representative of samples in the two gravity conditions. Images were acquired at varying magnifications to allow the visualisation of the features of interest. Scale bars are indicated in each figure, ranging between 2-20 µm.

The images demonstrate that the interaction of both the bacterium and the fungus with the meteorite material occurred under microgravity as well as terrestrial gravity. *S. desiccabilis* formed a contiguous biofilm in many areas of the rock’s surface in both the ISS and the Earth samples. Qualitatively, no gravity-dependent pattern for biofilm formation or cellular morphology was detected. *P. simplicissimum* successfully formed mycelia on the meteoritic rock fragments in both gravity conditions (Figure 4), with no evident qualitative morphological change. Similar results were observed for the Consortium samples, with the bacterium and the fungus interacting in a similar fashion in both gravity conditions and forming a mixed filamentous (*P. simplicissimum*) and rod-shaped cell (*S. desiccabilis*) biofilm on the rock surface. In all the biological samples, EDS spectra showed a tendency of both the bacterium (S5) and the fungus (S6), included in the Consortium (S7), to interact with minerals bearing principally magnesium, oxygen, silicon (silicates), and less frequently with iron, copper, sulphur, chromium, manganese and other metals, in both gravity conditions (S5-7). This is in accordance with the rock composition described above (Table1-2, Figure 2).

### 2.4 Microbial bioleaching of PGEs onboard the ISS

Measurement of the concentration of 44 elements in the liquid fraction was assessed by ICP-MS, as a measure of elemental dissolution from the meteorite rock (S8, Supplementary Excel file). Statistical analysis of the raw concentrations (ICP-MS data) by ANOVA revealed a p-value ≤ 0.05 for at least one of the two variables in analysis (*Gravity*, *Organism*), or their interaction, and then for biologically-relevant Tukey post hoc comparisons, for 22 elements out of these 44 (S9). Among these, PGEs captured our interest, and we focused our analysis on the three PGEs ruthenium (Ru), palladium (Pd) and platinum (Pt) (Figure 5, S10-11).

**Figure 5.**
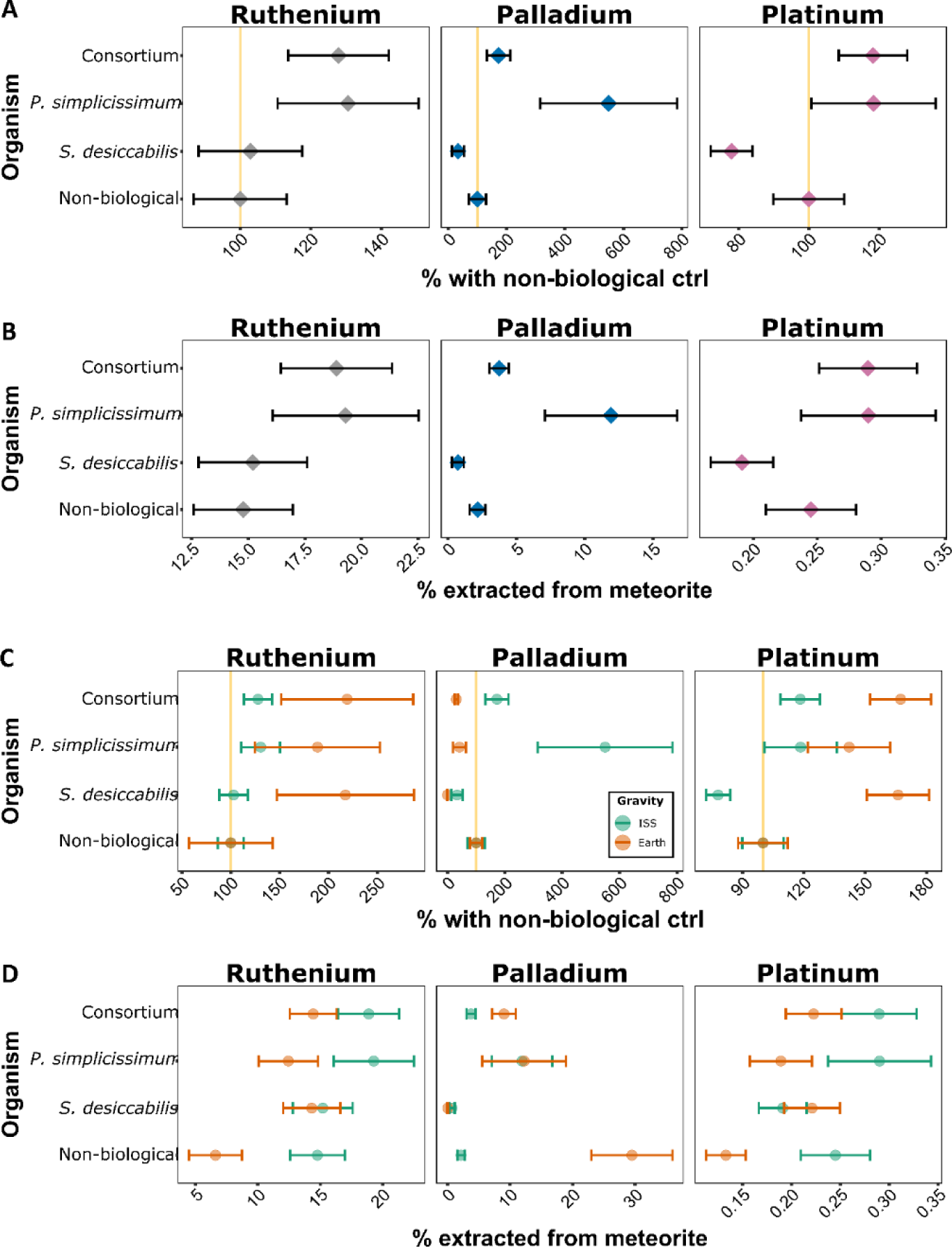
Platinum group element (PGEs) biomining. In panels A and B, data for the ISS experiment are shown. For each element, data are reported as: A) percentage differences with the non-biological control (%_NB_); B) percentage of total concentration in the meteorite (%_M_). For A and B, diamonds indicate mean values, error bars indicate standard error, yellow vertical lines in panel A indicate 100% values (i.e., same bioleaching efficiency as non-biological samples). For C and D, values are shown per element analysed, and per gravity condition (ISS = microgravity, green circles, Earth = 1 x *g*, orange circles). Data from the ISS and Earth experiment are shown C) as percentage differences with the non-biological control (%_NB_); D) as percentage of total concentration in the meteorite (%_M_). ISS data in C-D are the same as A-B. For C and D, circles indicate mean values; error bars indicate standard error; yellow vertical lines in panel C indicate 100% values (i.e., same bioleaching efficiency as non-biological samples).

Comparison by ANOVA (S9) suggests that the variable *organism* influences the dissolution of all the PGEs analysed. Focusing on the ISS samples only, p-values of Tukey pairwise comparisons for ruthenium, palladium and platinum were > 0.05 in all pairwise comparisons between biological and non-biological samples, between single species, and between single species versus the consortium (S10). Two exceptions were found for platinum, for *S. desiccabilis* versus *P. simplicissimum*, or versus the Consortium (p-values = 0.016 in both cases), suggesting different effect of the two organisms on platinum extraction (S9-10).

Metal concentration for all elements were normalised to the relative non-biological controls (%_NB_) for the ISS samples, to determine whether the organisms would enhance leaching under microgravity (Figure 5A, S11). When observing mean %_NB_ values to highlight general bioleaching trends in space, an enhanced bioleaching for PGEs was present in the presence of *P. simplicissimum* (ruthenium: 130.6±19.9%_NB_, palladium: 549.3±234.4%_NB_, platinum: 118.4±17.6%_NB_), while it was reduced or unaffected in the presence of *S. desiccabilis* alone (ruthenium: 102.9±14.7%_NB_, palladium: 33.6±20.1%_NB_, platinum: 78.0±5.9%_NB_). The Consortium %_NB_ were similar to those of *P. simplicissimum* alone (ruthenium: 127.8±14. %_NB_, platinum: 118.2±9.7%_NB_), except for palladium (Consortium: 172.4±40.2%_NB_ vs *P. simplicissimum*: 549.3±234.4%_NB_), whose %_NB_ values were between the fungus and the bacterium (Figure 5A, S11).

Elemental concentrations of ruthenium, palladium and platinum in the ISS samples were compared to those present in the meteorite (%_M_), to calculate the amount leached from the rock by non-biological and biotic processes under microgravity (Figure 5B, S11). Non-biological leaching released 14.8±2.2%_M_ ruthenium, 2.2±0.6%_M_ palladium, and 0.2±0.0%_M_ platinum present in the meteoritic rock. *P. simplicissimum* extracted 19.3±3.2%_M_ ruthenium, 11.9±4.8%_M_ palladium, and 0.3±0.1%_M_ platinum. *S. desiccabilis* extracted 15.2±2.4%_M_ ruthenium, 0.7±0.4%_M_ palladium, and 0.2±0.0%_M_ platinum. The Consortium leached 18.9±2.5%_M_ ruthenium, 3.7±0.7%_M_ palladium, and 0.3±0.0%_M_ platinum of the total in the meteoritic rock, respectively.

### 2.5 Microbial bioleaching of PGEs on Earth

Similarly to the ISS samples, we compared the Earth samples to determine whether the organisms would enhance PGEs leaching under terrestrial gravity (Figure 5C, S9-11). On Earth, p-values of raw concentration comparisons for ruthenium, palladium and platinum were ≤ 0.05 in all pairwise comparisons between biological and non-biological samples, with the exception of *P. simplicissimum* for ruthenium and platinum (S10), while all comparisons between biological samples had a p-value > 0.05.

Compared to non-biological samples (%_NB_), an enhanced bioleaching was observed in the presence of *S. desiccabilis* and *P. simplicissimum,* alone or in consortium, for ruthenium and platinum, with increases spanning between 142.2±19.9%_NB_ to 218.9±67.5%_NB_. In contrast, leaching of palladium was reduced on Earth in the presence of the microbial species (*S. desiccabilis*: 0.2±0.2%_NB_, *P. simplicissimum*: 41.6±22.5%_NB_, Consortium: 30.6±6.1%_NB_)(Figure 5C, S11). Relative to the amount present in the rock (%_M_), non-biological samples released 6.59±2.13%_M_ ruthenium, 29.47±6.47%_M_ palladium, and 0.13±0.02%_M_ platinum, *P. simplicissimum* extracted 12.4±2.4%_M_ ruthenium, 12.3±6.7%_M_ palladium, and 0.2±0.0%_M_ platinum. *S. desiccabilis* extracted 14.3±2.3%_M_ ruthenium, 0.1±0.1%_M_ palladium, and 0.2±0.0%_M_ platinum. Finally, the Consortium leached 14.4±1.9%_M_ ruthenium, 9.0±1.9%_M_ palladium, and 0.2±0.0%_M_ platinum (Figure 5D, S11).

### 2.6 Effect of microgravity on microbial-mediated PGEs bioleaching

Together with the influence of the *organism*, ANOVA results suggested that the variable *gravity* influences the dissolution of all the PGEs elements analysed (S9). The interaction between the variables *gravity* and *organism* has an effect for palladium and platinum, but not ruthenium (S9). To reveal the effect of gravity on overall leaching, we compared the raw concentrations of element extracted (ng/mL, equivalent to comparing %_M_) by performing Tukey pairwise comparisons between ISS and Earth samples harbouring the same organism(s). To analyse the effect of gravity on the organisms, we performed the same comparisons using a Student T test on the concentrations normalised for the non-biological controls (%_NB_), which allowed to remove the effect of abiotic leaching in our samples. For ruthenium and palladium, all comparisons between %_M_ and between %_NB_ produced p-values > 0.05 (S11, Figure 5C-D), with the exception of %_NB_ of the Consortium for palladium, which produced a p-value = 0.009 (ISS 172.4±40.2%_NB_ vs Earth 30.6±6.1%_NB_). For platinum, comparison between %_M_ for *P. simplicissimum* samples reported a p-value of 0.007, with leaching values of 0.3±0.0%_M_ (0.26±0.03ng/mL) on the ISS, and 0.2±0.0%_M_ (0.17±0.02ng/mL) on Earth (Figure 5C-D, S10-11). However, comparison between %_NB_ had p-value > 0.05. Comparison between %_NB_ for *S. desiccabilis* produced a p-value = 0.01 (ISS 78.0±5.9%_NB_ vs Earth 166.1±12.3%_NB_), while comparison of %_M_ was > 0.05. All other comparisons produced p-values > 0.05.

### 2.7 Bioleaching of other elements on ISS and Earth

Among the 22 elements analysed whose ANOVA revealed relevant results (see paragraph 2.4), and beside PGEs, 15 further elements produced p-values ≤ 0.05 for at least one biologically-relevant pairwise comparison, either when comparing the raw concentrations or the %_NB_ (S9, Supplementary excel file).

Comparing ISS samples only, pairwise comparisons for phosphorus revealed a p-value of 0.049 for *P. simplicissimum* versus non-biological control under microgravity, with 185.9±35.3%_NB_ and 0.26±0.04%_M_. Other pairwise comparisons for phosphorus and for the other 14 elements showed p-values > 0.05 (S9, S12-13, Supplementary excel file), nevertheless analysis of average extraction values allowed to highlight potential bioleaching trends. We chose a threshold of ≥1.5-fold change to compare mean %_NB_ values. Compared to non-biological samples, copper leaching was reduced (0.4-fold of the non-biological) by *S. desiccabilis*. while *P. simplicissimum* alone and in Consortium increased phosphorus leaching 1.6-fold (fungus) to 1.9-fold (Consortium). Moreover, the fungus (alone and in Consortium) compared to the bacterium increased bioleaching of phosphorus (fungus: 1.7-fold; Consortium: 1.6-fold), vanadium (fungus: 2.0-fold; Consortium: 1.8-fold) and copper (fungus: 2.8-fold; Consortium: 2.9-fold; S12-13). No average %_NB_ increase was present comparing the fungus and the Consortium (S12-13).

For Earth, p-values ≤ 0.05 for pairwise comparisons of biological versus non-biological samples (Supplementary excel file) were found for potassium (p=0.009 *S. desiccabilis*; p=0.005 Consortium), vanadium (p=0.015 *P. simplicissimum*; p=0.0007 Consortium), manganese (p=0.015 *S. desiccabilis*, p=0.048 *P. simplicissimum*, p=0.006 Consortium), iron (p=0.00006 *S. desiccabilis*; p=0.004 Consortium), nickel (p=0.004 *S. desiccabilis*; p=0.008 Consortium), strontium (p=0.042 *S. desiccabilis*; p=0.019 Consortium), zirconium (p=0.015 *S. desiccabilis*; p=0.006 Consortium), molybdenum (p=0.003 *S. desiccabilis*), barium (p=0.0016 Consortium) and europium (p=0.033 Consortium). When comparing average %_NB_ as above (≥1.5-fold increase threshold), *S. desiccabilis* showed a higher extraction capacity than *P. simplicissimum* for 3 out of 15 elements, namely iron (1.7-fold), cobalt (1.9-fold), and molybdenum (1.6-fold), while the fungus compared to the bacterium increased copper leaching (2.1-fold). The fungus also increased copper extraction compared to the Consortium (2.4-fold), while this latter did not improve bioleaching compared to the single organisms (S12-13, Supplementary excel file).

To highlight the effect of gravity on bioleaching, comparisons between gravities in same-organism samples were analysed (S12-13, Supplementary excel file). P-values of raw concentration pairwise comparisons were ≤0.05 only for lutetium with the fungus alone (0.057±0.001%_M_ ISS vs 0.003±0.001%_M_ Earth). When comparing %_NB_, p-values of the Consortium were ≤0.05 for sodium (117.57±6.24%_NB_ ISS vs 86.53±3.78%_NB_ Earth), copper (125.38±13.51%_NB_ ISS vs 31.92±14.42%_NB_ Earth) and zinc (134.27±17.05%_NB_ ISS vs 92.53±4.76%_NB_ Earth). Zinc is in addition to the 15 elements discussed above, since raw concentration pairwise comparisons did not report relevant p-values (S9), while pairwise comparisons of the %_NB_ for the Consortium produced a p-value = 0.04 (Supplementary excel file). All other pairwise comparisons, either for the raw concentrations or for the %_NB_, were > 0.05. Similarly to above, we analysed mean %_NB_ values to highlight general trends related to the effect of gravity. *S. desiccabilis* leaching showed ≥1.5-fold increase leaching on Earth compared to ISS for 10 out of 15 elements, namely barium (2.0-fold), cobalt (2.3-fold), europium (2.1-fold), iron (4.1-fold), manganese (2.0-fold), molybdenum (2.9-fold), nickel (2.0-fold), strontium (1.7-fold), vanadium (4.1-fold) and zirconium (1.9-fold). *P. simplicissimum*, alone and in Consortium, showed higher average %_NB_ in microgravity compared to Earth gravity for phosphorus (fungus: 1.7-fold; Consortium: 1.6-fold) and copper (fungus: 1.6-fold; Consortium: 3.9-fold), but higher on Earth for 9 out of 15 elements, namely barium (fungus: 2.3-fold; Consortium: 2.9-fold), cobalt (fungus: 1.5-fold; Consortium: 2.2-fold), europium (fungus: 1.6-fold; Consortium: 2.1-fold), iron (fungus: 2.5-fold; Consortium: 3.6-fold), manganese (fungus: 1.8-fold; Consortium: 2.0-fold), molybdenum (fungus: 1.8-fold; Consortium: 2.4-fold), nickel (fungus: 1.5-fold; Consortium: 1.9-fold), strontium (fungus: 1.6-fold; Consortium: 1.8-fold) and vanadium (fungus: 2.3-fold; Consortium: 3.3-fold).

### 2.8 Effect of microgravity on abiotic leaching

To test the effect of the gravity condition on the abiotic leaching from the meteorite rock, comparisons between non-biological samples in microgravity (ISS) and Earth gravity (Earth) were measured.

For PGEs (Figure 5C-D, S10-11), this comparison reported a p-value ≤ 0.05 for palladium and platinum, but not ruthenium (S10-S11), with mean extraction from the rock of 14.8±2.2%_M_ in microgravity versus 6.6±2.1%_M_ under terrestrial gravity for ruthenium, 2.2±0.6% _M_ on ISS versus 29.5±6.7% _M_ on Earth for palladium, and 0.2±0.0% _M_ in microgravity versus 0.13±0.02% _M_ under terrestrial gravity for platinum.

Besides palladium and platinum, other 9 elements reported a p-value ≤ 0.05 for pairwise comparisons of non-biological controls under microgravity versus Earth gravity. These are, in order of atomic number, sodium, aluminium, scandium, iron, cobalt, nickel, strontium, molybdenum, and erbium (S9, Supplementary excel file).

Compared to Earth samples, enhanced abiotic leaching in microgravity was observed for aluminium (6.8-fold increase in ISS vs Earth; ISS: 1.52±0.95% _M_, Earth: 0.22±0.12% _M_), scandium (3.4-fold; concentration in the rock was not detected. Absolute values in the leachate: ISS = 0.020±0.008 ng/mL, Earth = 0.006±0.002 ng/mL), iron (4.3-fold; ISS: 0.12±0.01%_M_, Earth 0.03±0.01%_M_), cobalt (2.5-fold; ISS: 0.50±0.09%_M_, Earth: 0.21±0.07%_M_), nickel (2.4-fold; ISS: 0.15±0.01%_M_, Earth: 0.06±0.01%_M_), strontium (1.8-fold, ISS: 0.39±0.01%_M_, Earth: 0.21±0.04%_M_), molybdenum (2.9-fold, ISS: 0.03±0.01%_M_, Earth: 0.01±0.00%_M_) and erbium (2.4-fold, ISS: 0.003±0.0004%_M_, Earth: 0.001±0.0004%_M_). Abiotic leaching was higher under terrestrial gravity for sodium (1.4-fold increase on Earth; ISS: 0.57±0.07%_M_, Earth: 0.82±0.11%_M_). Silicon showed a similar result (ISS: 0.0097±0.001%_M_, Earth: 0.011±0.001%_M_), but these values might be suboptimal due to the compromised silicon detection in the rock (see materials and methods), and hence excluded from further analysis (S9, Supplementary excel file).

### 2.9 Metabolomics of biomining-related features

In order to determine whether microgravity alters the metabolome of biomining microorganisms, and thus plays a role in their different behaviour in space, we conducted a metabolomics analysis of the liquid fraction of the samples after the experiment. Principal component analysis (PCA) of the whole set of samples, both in microgravity (ISS) and Earth gravity (Earth), showed an overlapping of all the samples, with the exception of a single outlier (an ISS *S. desiccabilis* sample), indicating the lack of a significant component that could allow the discrimination of the samples based on gravity condition or organism (Figure 6A). PC1 and PC2 taken together explain only a 60.5% of the observed variance. The PCA analysis is in accordance with the volcano plot analysis (S14), showing a larger number of up- or downregulations of features in space compared to Earth for the fungus-containing samples, compared to those non containing the fungus.

**Figure 6.**
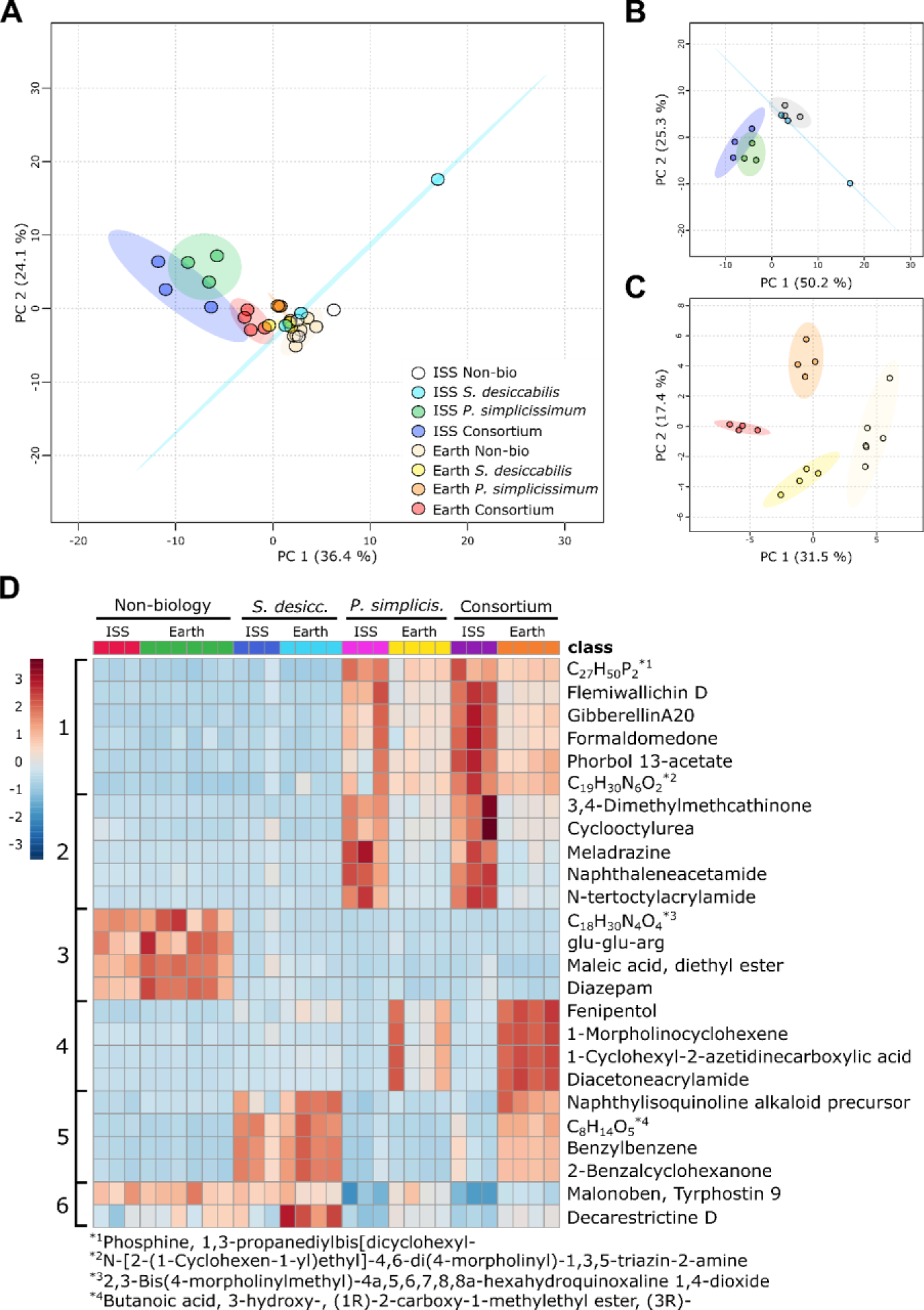
Metabolomic analysis of microorganisms during microbe-meteorite interaction in space and on Earth. A) Principal component analysis (PCA) representing all the samples; B) PCA showing ISS samples only; C) PCA showing Earth samples only. Ellipses correspond to 95% confidence intervals. D) Heatmap showing the top 25 features measured in the metabolomics analysis, clustered by gravity condition and organism. Each column represents a single sample. Numbers of the left represent visual clusters suggesting metabolomic patterns dependent on the organism and/or the gravity condition. Heatmap legend is on the left side of the figure, indicating red colours for higher concentrations and blue for lower concentrations of features in each specific sample. Features with long names were indicated with their chemical formula, with asterisks showing the complete names at the bottom of the figure.

To better appreciate metabolomics characteristics regardless of the gravity condition, PCAs of space only (Figure 6B) and Earth only (Figure 6C) sample were produced. PCA analysis of the space samples (Figure 6B) shows *S. desiccabilis* cluster partially overlapping with the non-biological cluster, with the exception of one outlier sample. It also shows the fungus-containing samples clustering together and separately from the non-biological controls and the *S. desiccabilis* samples. PC1 and PC2 together explain a total of the 75.5% of the variance.

PCA analysis of the Earth samples alone (Figure 6C) shows 4 separate clusters, indicating that, under terrestrial gravity conditions, *S. desiccabilis*, *P. simplicissimum* and the two organisms in the Consortium present different metabolic profiles. Notably, the separation in four distinguished clusters may indicate a minor or null effect of the contaminants (S3), if present, on the metabolomics analysis.

Figure 6D shows a heatmap of the top 25 features measured in the liquid fractions of the BioAsteroid samples. The heatmap allows the identification of clear clustering of features, specific to the gravity condition and/or the organism. Cluster 1 represents metabolites produced in the presence of the fungus, either as a single culture or in the Consortium, in both gravity conditions. A higher presence in microgravity (ISS) is however evident. Cluster 2 represents features expressed almost solely in the presence of *P. simplicissimum* in space. Cluster 3 shows features present in the non-biological control samples regardless of gravity condition. Complementary to cluster 2, cluster 4 shows features associated with the presence of the fungus when subjected to Earth gravity. Cluster 5 shows features mostly associated with the presence of *S. desiccabilis*, regardless of the gravity condition when alone, and under terrestrial gravity when in Consortium. This might indicate either lack of production of these features in the presence of the fungus, or their fungal degradation/metabolization, in microgravity. Cluster 6 represents features less expressed in the presence of the fungus in both gravity conditions. The patterns shown here are similar to those observed in the heatmap showing the top 70 features (S15).

## 3. Discussion and conclusion

Space biomining is an area of space activity with substantial interest, since it offers the promise of supporting *in-situ* resource utilisation (ISRU), obtaining and recycling materials in extraterrestrial settlements, and enhancing the self-sustainability of human space exploration^4,16^. In this work, we tested the proof of concept for bacterial and fungal bioleaching of asteroidal rock material (L-chondrite) in microgravity, onboard the ISS.

Focusing on PGEs, we observed that the mean leaching of palladium for the fungus *P. simplicissimum* was 549.3% of the non-biological control on the ISS, and mean leaching of ruthenium and platinum was slightly higher than the non-biological control. These results suggest that a fungus can be used to increase mean leaching rates of certain PGEs from asteroidal material in microgravity. However, we also observed that there was great variability around the mean, leading to p-values > 0.05 when biological extraction was compared to non-biological controls. This variability could be caused by intrinsic differences in growth rates between our replicates, but it might also reflect genuine heterogeneities in biomining. As we showed in our geological and geochemical analysis, the meteorite used is a heterogeneous mixture of mineral fragments which, despite being quite consistent in concentration, could have been exposed on the surface in different proportions. Thus, variability could be an intrinsic part of biomining process in this experiment, especially since we were working with small volumes of rock pulp and with a small number of replicates, an inevitable limitation of space experiments.

We observed different results with the bacterium *S. desiccabilis* in microgravity. PGEs mean leaching was similar (ruthenium) or lower (palladium and platinum) with the organism compared to the non-biological control, suggesting that this organism did not successfully extract PGEs from L-chondrite in microgravity. Although biofilms are generally thought to be beneficial for bioleaching^24,48^, they have also been shown to protect metal and concrete surfaces from biotic and abiotic corrosion^49–51^, through the production of polysaccharides. Hence, a possible explanation of the reduction in mean leaching for *S. desiccabilis* in microgravity can reside in its well-known biofilm and polysaccharide production capacity^52^. SEM images show biofilm formation on the rock surfaces in both gravity conditions, although their sporadic nature does not allow us to definitely link biofilm growth to potential leaching rates in the two gravity conditions tested. The general impairment of *S. desiccabilis* bioleaching in microgravity compared to observations on Earth is partially in accordance to our previous space biomining experiment, BioRock^31,32^, in which *S. desiccabilis* extraction of REEs and vanadium from basaltic rock (a different rock from that of this experiment) had a reduced trend in microgravity, compared to simulated Martian and terrestrial gravities. In BioRock, *S. desiccabilis* extracted a mean value of vanadium of 184.92% of the non-biological control from the basalt rock in microgravity^31^, compared to 71.40% from the L-chondrite in this experiment, suggesting an effect of the starting rock material. Only the REEs europium and lutetium were bioleached by the bacterium in this work (mean values), compared to the whole range of REEs extracted in BioRock^32^. Lutetium is a heavy REEs, which were preferentially extracted in BioRock and in other experiments reported in the literature^32,53^.

When we used a consortium of both the bacterium and fungus together, we found that mean leaching rates were higher than non-biological controls for all three PGEs in microgravity, but again, variability in mining efficiency meant that p-values were greater than 0.05. Mean values reported a similar enhancement between the consortium and the fungus alone for ruthenium and platinum, suggesting the fungus dominates the bioleaching activities for these elements, while for palladium we did not observe the same enhancement with the consortium as with the fungus alone. This may be explained by the fact that the bacterium strongly reduced mean palladium leaching when used alone (33.6%_NB_) and may counteract the beneficial effects of the fungus in this case.

We also investigated the bioleaching on a range of 41 further elements, 15 of which showed alterations between the conditions tested. Aside from palladium, total fungal leaching of phosphorus (compared to non-biological) was enhanced in microgravity, showing how microorganisms might be employed to extract other crucial elements required in industry, and that materials for life support systems might be bioextracted from asteroidal materials. All other comparisons between ISS samples provided p-values > 0.05. However, mean values for copper were lower in the presence of the microorganisms, particularly the bacterium, compared to the non-biological control in microgravity. This suggest that, in some cases, abiotic leaching could be a preferable way to mine in space, with organisms potentially inhibiting leaching, as we also observed for *S. desiccabilis* and the PGEs. The fungus, alone and in consortium, caused higher mean %_NB_ bioleaching of phosphorus compared to the non-biological control, and of phosphorus, vanadium and copper, compared to the bacterium. Similarly to ruthenium and platinum, mean values for the fungus and the consortium were similar.

Underlying our biological observations may be changes in the abiotic rate of leaching. Compared to terrestrial gravity controls, 11 elements (including 2 PGEs) showed changes in their abiotic leaching in microgravity. Two of these showed reduced leaching under microgravity, while the remaining 9 showed an increased leaching, including platinum (1.84-fold increase in ISS vs Earth). This is consistent with our previous observation that mean abiotic leaching values for REEs increased with decreasing gravity regimens (Earth gravity, Mars gravity and microgravity, respectively) onboard the ISS^32^. Among those that showed decreased abiotic leaching under microgravity was the PGE element palladium (13.6-fold decrease in ISS vs Earth), where absolute leaching rates for *P. simplicissimum* were similar for Earth and space. This means that the fungus allowed a net 13.2-fold increase in palladium extraction in space, opening fascinating scenarios for future space biomining and ISRU technologies. Similar results were found for sodium. A reduction in abiotic leaching of some elements in microgravity is difficult to explain. One possible hypothesis is that the different fluid dynamics in microgravity (i.e. lack of convection), compared to terrestrial gravity, could have enhanced local saturation of the leached elements around the rock surface^54^, in turn reducing abiotic leaching. However, this would not explain why this was only observed for certain elements. These results emphasise the necessity of selecting the most appropriate technology when planning ISRU. Indeed, they demonstrate that biomining may be beneficial for palladium and sodium (and probably silicon) extraction in space, for instance, but not for other elements. They also indicate that the benefits of utilising microorganisms in space mining lie not only in their ability to increase absolute leaching in some cases, but also to maintain stable leaching rates against a potential reduction in abiotic leaching in other instances.

When analysing the Earth results, many comparisons between biological and non-biological controls revealed a p-value ≤ 0.05. The presence of *S. desiccabilis*, alone or in consortium, had a positive effect on bioleaching of ruthenium, platinum, potassium, vanadium, manganese, iron, nickel, strontium, zirconium, and molybdenum. *P. simplicissimum*, alone and in consortium, improved the extraction of vanadium and manganese, compared to the non-biological control. The consortium alone improved the extraction of barium and europium, where the single species did not enhance leaching in comparison to the non-biological control. When comparing organisms in the terrestrial samples, *S. desiccabilis* showed higher mean leaching capacity (%_NB_) for 5 of the 18 (considering the three PGEs) elements compared to the fungus. *Viceversa*, the fungus extracted only copper better than the bacterium and the consortium, and the consortium did not improve bioleaching compared to single species. Taken together, these results indicate *S. desiccabilis* has a wider bioleaching capacity than *P. simplicissimum*, in terms of number of elements extracted from the meteorite materials, under terrestrial gravity.

We therefore compared experiments in microgravity and in terrestrial gravity for all the elements we studied, to highlight the effect of gravity on the bioleaching efficacy of the two organisms tested. Aside from platinum, *P. simplicissimum* %_M_ extraction of lutetium, and consortium %_NB_ extraction of sodium, copper and zinc, were higher in microgravity than terrestrial gravity. All other comparisons provided a p-value > 0.05, however mean %_NB_ bioleaching values of 10 elements for the bacterium, and 9 for the fungus and the consortium, were lower in microgravity compared to terrestrial gravity.

Taken together, these results show the complexity of the biomining process, particularly under space conditions. Understanding the effect of microgravity on bioleaching will depend on the microorganism, whether it is alone or in consortium, and on the characteristic of the element/mineral/rock, or a combination of these. Microgravity can also influence abiotic leaching. These results highlight the need to carefully select optimal combinations of microorganism(s), rock substrate, and leaching conditions for successful biomining setting, either in space or on Earth. It also indicates that it is difficult to predict *a priori* how to go about extracting a chosen element from asteroids, without carrying out experiments to test each condition. In some instances, abiotic leaching should be preferred to bioleaching, and costs/benefits should be carefully considered case by case.

Comparisons of the biological rates of leaching on Earth versus space needs to be tempered with caution, due to the contaminations observed during the CFU analysis for some of the biological Earth samples. The analysis of the DNA extracted from the *S. desiccabilis* and the Consortium Earth samples suggests that either the contamination was not present in the original samples, or that *S. desiccabilis* dominates it, in accordance with the CFU numbers (i.e., contaminants’ CFU concentration is 5-6 orders of magnitude lower than that of *S. desiccabilis*, S3). Results on these samples are therefore considered accurate. Results on the fungal Earth samples suggested a contamination from the order Bacillales (probably belonging to the genera *Bacillus* and/or *Paenibacillus*). We can hypothesise that any relevant effect of the contamination would have likely influenced all the affected samples equally. From the metabolomics analysis we note this is not the case, as suggested by the absence of contaminant-related clusters in the heatmap results and the PCA analysis. This might indicate that, if any effect was present, this does not have a relevant impairing role on the results. We were not able to repeat the experiment to test these hypotheses, due to the unavailability of identical material from the same L-chondrite meteorite. However, even when the Earth fungal results require these caveats, the other results are completely valid.

We were also interested to determine whether microgravity conditions would alter the metabolome of biomining organisms. Metabolomics analysis revealed clear biologically relevant patterns that highlight the influence of the space condition and of gravity on the microbial metabolism. PCA analysis on the ISS samples reveals two main clusters, partially overlapping, namely one that groups the non-biological controls with the bacterium samples, and a second that groups together the fungus-harbouring samples. Indeed, of the biological systems we studied, the fungal metabolic pattern was the most influenced by microgravity, both in the consortium and as a single-species (Figure 6D, Clusters 1-2; S15 Cluster 1). This potential effect of microgravity on the secondary metabolite production of the fungus, only partially influenced by its presence in a consortium or in a single species culture, is corroborated by the PCA clustering.

Data on microbial metabolomics under real or simulated microgravity are still scarce and are mostly focused on bacterial rather than fungal metabolites^55^. Secondary metabolites of interest in previous experiments were mostly antibiotics, but also molecules such as poly-β-hydroxybutyrate (a polyester)^56^. Their production has been seen to increase, decrease or be unaffected by the gravity conditions, indicating experimental methods should be carefully selected in order to have comparable results, but also a non-trivial effect of the space conditions on microbial metabolism^55^.

Analysis of the single metabolites allowed insights into the metabolic pathways associated with the presence of the meteorite, the bioleaching capacity of the organism and the gravity condition. Specific mechanisms of *Sphingomonas* spp. and *P. simplicissimum* bioleaching include organic acids production, such as citric, oxalic, malic and glucuronic acids^57,58^. While we could not identify these three specific organic acids in our result, features of interest from the bioleaching perspective include carboxylic acids and siderophore-associated molecules. Linked with the presence of *S. desiccabilis* alone or in the consortium in both gravities: (i) a butanoic acid derivate, (ii) the carboxylic acid 2’-Deoxymugineicacid, (iii) Quinoline-3,4-diol linked with biofilm formation. In fungal ISS samples we identified (i) Phosphine, 1,3-propanediylbis[dicyclohexyl-, which has been associated with chemical palladium-catalysed carbonylation (although its biological activity is unknown)^59^, (ii) the fatty acid Undeca-2,5-dienal, and (iii) 2,3-Dihydro-2,3-dihydroxybenzoic, both acid associated with bacterial siderophore biosynthesis. Linked with the fungal presence on Earth samples we found (i) 1-Morpholinocyclohexene which, although not reported to be associated to bioleaching *per se*, belongs to the morpholino-compounds class which are known to perform abiotic metal dissolution^60^, (ii) the carboxylic acids 1-Cyclohexyl-2-azetidinecarboxylic acid, (iii) the carboxylic acids 1-Cyclohexyl-2-azetidinecarboxylic acid and (iv) hexanoic acid. These features are thus good candidate for future research, focused on both understanding the bioleaching mechanisms of the two organisms, and bioengineering endeavours.

Notably, a number of compounds of pharmaceutical interest were also identified. These include Lastar A^61^, Flemiwallichin D (a chalcone flavonoid)^62^, Sarcoehrendin D (a prostaglandine derivative), and laurolactam, all associated with the presence of *P. simplicissimum* in ISS samples (S15, Cluster 1). Laurolactam is used for the production of plastics such as polyamide 12 (nylon 12) and copolyamides^63^. Although its natural bioproduction has not been reported, its presence in the fungal-containing cluster opens speculation on potential bioplastic production in space. Many of the features present in the top 70 list (S15) were of unknown relevance for the aim of our experiment, showing that we still have much to learn about the complex metabolic restructuring that occurs under space conditions as well as the functions of many metabolites. Compared to space samples, PCA analysis of the Earth samples forms four diverse groups. These results suggest that under terrestrial gravity conditions the metabolome is more plastic and adaptable, allowing diverse response to different biological conditions, but note our observations about contaminant species.

These data show that microgravity can have profound effects on biomining organisms. Although we have identified a range of metabolites that might be linked to biomining, the role of many of these metabolic changes remains enigmatic. This demonstrate that understanding and predicting biomining processes on asteroids may ultimately be linked to our grasp of the metabolic response of organisms in space. In particular, as we show here, changes in the metabolome of biomining organisms may even lead to changes in other compounds with human health and industrial implications.

We carried out investigations on the cell numbers using different methods (Figure 3, S2-3). These results provide different answers depending on the technique used. For instance, the higher final cell concentration for *S. desiccabilis* measured with the CFU method is not corroborated by flow cytometry, which did not show differences between gravity conditions. CFU results for *P. simplicissimum* identified a higher concentration under microgravity compared to terrestrial gravity, while a lower concentration is reported by the other two methods. There results may have some bearing on the widely inconsistent reports of the effects of space on microorganisms ^55,64^. The interplay of viability, metabolic states and other factors may influence how the effects of microgravity are manifested in cell numbers depending on the method used to measure them. Thus, experiments investigating the effects of microgravity should choose the method of analysis carefully and preferably more than one method should be used.

Finally, we provided a demonstration of bacterial and fungal interaction with extraterrestrial minerals under microgravity conditions, on the ISS. Both bacterial biofilms and fungal mycelium grew on the meteorite fragments, both alone and in consortium. Qualitatively speaking, the microorganisms are frequently associated with magnesium-, oxygen- and silicon-harbouring minerals (silicate minerals), rather than sulphur- and iron-ones (sulphide minerals). This could plausibly be caused by the greater diversity of elements available in the silicate minerals compared to the sulphides, which are relatively pure. No particular morphological difference could be reported when observing microbial-mineral interaction in microgravity or on Earth. Although our results are of qualitative nature, to our knowledge they represent the first demonstration of metabolically-active microbe-meteorite interaction, and of mycelium formation on extraterrestrial materials, in space. The results reported here are relevant not only from the industrial perspective of biomining, either space or Earth associated, but because rock weathering can be used to produce soils and release elements for life support systems^2,4–6,65^, such as phosphorus, potassium and iron.

## 4. Material and methods

### 4.1 Strains and medium

The microbial species used for this work were the bacterium *Sphingomonas desiccabilis* CP1D ^66^, a Gram-negative, non-motile and non-spore-forming bacterium, first isolated from soil crusts in the Colorado plateau ^39^ with demonstrated capacity to extract metals during spaceflight ^31,32^, and the fungus *Penicillium simplicissimum* DSM 1078 (DSMZ), an Ascomycota known for its capacity to perform biomining ^43,67^.

The medium used for this experiment was a solution of 50 % (v/v) R2A medium (Reasoner & Geldreich, 1985), chosen to encourage bacteria to extract nutrients from the meteorite. Five millilitre of medium were used in this experiment for each sample. The medium composition was (g L^-1^): yeast extract 0.25; peptone 0.25; casamino acids 0.25; glucose 0.25, soluble starch 0.25, Na-pyruvate 0.15; K_2_HPO_4_ 0.15; MgSO_4_.7H_2_O 0.025 at pH 7.2.

The fixative selected to prevent degradation of the biological portions and to stop microbial growth after the end of the experiment was RNAlater (Thermo Fisher), an aqueous and non-toxic storage solution compatible with the astronaut safety requirements on the ISS. One millilitre of fixative was used for each sample, with a final volume ratio of 1:5 fixative-medium.

### 4.2 Rock samples

The extraterrestrial rock sample used for this work was the Northwest Africa (NWA) 869 meteorite, a L3-6 chondrite regolith breccia^68,69^. A portion of the meteorite was crushed into irregular pieces of approximately 1-3.5 mm of diameter. Rock fragments were aliquoted in samples of 0.79±0.14 g (mean±st. dev.) each and sterilised by dry-heat sterilisation in a hot air oven (Carbolite Type 301, UK) for 4 h at 250 °C. The meteorite was characterized as described in sections 4.7, 4.8 and 4.10.

Average surface area of the meteorite fragments was measured by gas adsorption analysis (Quantachrome, Nova Touch), to measure the surface available to the microorganisms for bioleaching and biofilm formation. The average surface area for the meteorite’s fragments was 1.941±0.181 m²/g (mean±standard error, n=3), hence each sample provided ∼1.5 m^2^ of surface available.

### 4.3 Sample preparation for the spaceflight

Single strain cultures of each species were desiccated on sterile rock samples.

For *S. desiccabilis* CP1D, an overnight culture of the microorganism was grown in 100% (v/v) R2A medium at 20-22 °C. When the culture reached stationary phase (OD_600_ = 0.88±0.09, corresponding to 4.7 x 10^10^ CFU/mL), crushed sterile meteorite was soaked in 1 mL of the bacterial culture for *S. desiccabilis* and Consortium cultures and samples were air-dried at ∼20-25 °C in a laminar flow hood under sterile conditions.

The mycelium of a 7-day old pre-culture of *P. simplicissimum* (50 mL) was dissolved by sonication (Microson ultrasonic cell disruptor, Misonix) with continuous pulse at setting 3 for 2 minutes, and then filtered through a sterile cotton bud to remove larger bids of mycelium and obtain a homogeneous fungal solution. This procedure did not alter fungal viability (data not shown). One mL (containing ∼6 x 10^6^ CFU/mL) of the resulting liquid fraction was used to inoculate the sterile crushed meteorite samples *P. simplicissimum* and Consortium in sterile 6-well plates, and these were air-dried overnight at ∼20–25 °C with a sterile procedure within a laminar flow-hood.

Non-biological controls were sterile crushed meteorite samples without cell inoculation.

After preparation, all samples were stored at room temperature (∼20-25 °C) until integration in the BioMining Reactors (BMRs).

### 4.4 Flight experimental setup

Flow diagram summarising the BioAsteroid experiment setup is available in Figure 1C. Sample, medium and fixative integration into each Experiment Unit (EU)^70^ (KEU-RK, from Kayser Italia, http://www.bioreactorexpress.com/) was performed under aseptic conditions before the launch. Each EU was composed of two BioMining Reactors (BMRs), which are culture chambers of 15 x 14 x 23.2 mm, that can contain 6 mL of liquid volume after hardware activation and medium injection. After integration, the culture chamber is delimited by the meteorite fragments, allocated on an aluminium grid to avoid dispersion of the rock pieces in the culture chamber, on one side, and a semipermeable silicone rubber membrane, to allow gas diffusion, on the remaining five sides. A small sterile piece of cotton ball was inserted between each rock sample and the EU back cover, to protect the rock pieces from excessive shaking during the rocket launch and space operations. Each BMR is connected to a 5 mL medium reservoir and a 1 mL fixative reservoir that were activated at the appropriate time. Each EU was integrated in an Experiment Container (EC, KIC-SLA-E3W, Kayser Italia) featuring semipermeable membrane for gas exchange and a transparent window that allows the direct observation of the experiment. ECs were equipped with temperature loggers (installed in one EUs; Signatrol SL52T sensors, Signatrol, UK) and accelerometers (in all EUs on ISS). A complete description of the EU can be found in ^70^. A total of 12 samples in 6 EUs for the flight experiment and 18 samples in 9 EUs for the Earth samples were prepared on different timelines. After integration of the 6 flight EUs, occurred between September 29^th^ and October 2^nd^, 2020, they were shipped to NASA Kennedy Space Centre (Florida, USA), while being stored at room temperature, and launched to the International Space Station (ISS) on a SpaceX Falcon-9 rocket (Commercial, Resupply Mission 21 mission) on December 6th, 2020. On arrival to the ISS, the samples were stored at room temperature (23.0 °C, temperature loggers) until installation into the microgravity (non-centrifuged) slots within the two KUBIK (ESA) incubators (5 and 6, Figure 1B) aboard the ISS, previously set to 20 °C, on December 20^th^, 2020, when the automatic timeline of the EUs was activated and medium (5 mL) was injected in consecutive manner to each culture chamber. All crew activities were performed by NASA astronaut Michael S. Hopkins. Samples grew for 19 days at 19.5 °C (temperature loggers). At the end of the experiment, 1 mL of fixative was automatically injected into the culture chambers on January 8^th^, 2021) and hardware were cold stored at 1.5-11.5 °C (logged data). On orbit, the EUs were stored in the MELFI hardware, and were downloaded to Earth in cold storage bags (NASA-supplied passive temperature controlled facilities), in the SpaceX CRS-21 Dragon capsule (the same vehicle as for upload). Samples were shipped in cold storage to the University of Edinburgh (UK), where samples were retrieved after 12 days from the fixative injection.

Of the 9 EUs containing 18 Earth samples, 2 EUs, containing 4 non-biological controls, were prepared on March 16^th^, 2021, while the remaining 14 EUs were integrated on April 19^th^, 2021. All the Earth samples were subject to analogous procedures and conditions to those occurring in the flight hardware, with incubation at 20 °C in a laboratory incubator (Memmert). Medium and fixative were injected by manual manipulation of the appropriate screws. Fixative injection occurred after 19 days from the medium injection, similarly to the flight experiment. After fixative injection, EUs were stored at 8 °C for 12 (samples prepared in March) or 14 (samples prepared in April) days, until sample retrieval. The difference in storage timing was due to technical reasons and did not affect the results (data not shown).

### 4.5 Post-flight sample recovery

Samples were recovered separating the culture liquids, the meteorite fragments, the metal grids and the membranes.

Liquid cultures were treated differently depending on the species. While non-biological controls and *S. desiccabilis* samples did not require pre-treatments, liquid samples containing the fungus were homogenised by sonication (Microson ultrasonic cell disruptor, Misonix) with continuous pulse at setting 3 for 60 sec and then filtered through a sterile commercial cotton ball, to dissolve the mycelium and obtain a homogeneous solution. An aliquot of 1 mL was recovered from each liquid sample and immediately stored at -80C for the metabolomics analysis, 0.5 mL were collected for pH measurement, 0.2 mL were collected for CFU and optical density analysis. 0.25 mL were collected for flow cytometry analysis and treated as described below, the remaining aliquot of the liquid cultures were treated for ICP-MS analysis as described in the dedicated section.

An aliquot of the rock fragments was recovered, washed once with sterile water and air-dried at ∼20-25 °C in a laminar flow hood under sterile conditions. These samples were analysed by XRD (data not shown) and Raman. Other representative aliquots of rock fragment were stored in 4% (v/v) formaldehyde at 4°C to preserve biofilms and mycelia. Remaining rock fragments were treated for scanning electron microscopy analysis as described below. This latter process was also performed for the membranes, metal grid and cotton samples.

### 4.6 Final cell concentration

Final cell concentration was measured from the liquid fraction of the samples using three distinct methods: (i) measurement of the turbidity of the culture by spectrophotometric analysis; (ii) flow cytometry; (iii) counts of colony forming unit (CFU).

(i) Optical density (OD) was measured at wavelengths (λ) of 600nm and 530nm from 100 µL of the liquid fraction of each sample. Traditional OD_600_ was used to measure final cell concentration. However, due to the iron bioleaching from the L-chondrite, the liquid fraction had a strong orange/red coloration for some samples, which influenced the measurement. For this reason, and for the presence of the fungus^47^, non-standard OD_530_ was also measured (S2).

(ii) Flow cytometry was measured with a LSR Fortessa machine (BD Biosciences). Equipped with a 405nm laser, detecting the emissions of Calcofluor White Stain (fungi-specific dye, Sigma Aldrich) binding through a 450/50nm band pass filter, and a 488nm laser with a 530/30nm filter to excite BacLight Green Bacterial Stain (bacteria-specific dye, Invitrogen), as were forward (FSC) and side (SSC) scattering. A volume of 250 µL of the liquid fraction of each sample was washed once with a filter-sterile solution of Tween 80 at 0.1% (v/v) in PBS, then cells were fixed for 15 min at room temperature in a filter-sterile solution of 1% (v/v) formaldehyde in PBS. Finally, the liquid was removed and replaced after centrifugation for 5 minutes at max speed, with 250 µL of filter-sterile PBS. Samples were stored at 4°C until analysis. To have an estimation of final cell concentration, a volume of 100 µL of the liquid fraction of the samples, appropriately diluted in the diluent (PBS filtered with a 0.22 µm nylon filter), were acquired at a flow rate of 2-3 µL/sec, and all events were counted. Samples were stained with BacLight Green at a final concentration of 0.1 µM, Calcofluor White at a final concentration of 0.25 µg/mL, both or none. When possible, each sample was measured twice per dye (i.e., 2 technical replicates). Appropriate gating was constructed using the software BD FacsDiva 8.0.1, to distinguish bacterial from fungal cells (gating strategy is reported in S16). Events in Bacteria and Fungi gates were counted and considered as single cells, to reconstruct final cell concentrations, expressed as cells/mL.

(iii) To measure colony forming units (CFU), serial dilutions of the liquid fraction were prepared, and 6-10 spots of 10 µL (for a total volume of 60-100 µL, respectively) of each dilution were spotted on R2A solid medium. These were incubated at room temperature for 2-5 days, until single colonies became visible. These were counted from the lowest dilution in which they were clearly distinguishable, and colonies of each spot, for each sample, were summed. Final CFU concentration (CFU/mL) was then calculated with the formula: [(total colonies) x dilution) / total volume].

The 3 methods described above were compared building a growth curve for each organism (*S. desiccabilis* or *P. simplicissimum*) for 19 days (the timeframe of this experiment), and measuring cell concentration at each datapoint with the 3 techniques (S17-18).

As CFU showed a potential bacterial contamination of some of the samples (three of three *P. simplicissimum* ISS samples, two of four *S. desiccabilis* Earth samples, four of four *P. simplicissimum* Earth samples, three of four Consortium Earth samples; S3), the genomic DNA of the affected samples, as well as that of the isolated contaminant species, was extracted with DNeasy PowerLyzer Microbial Kit (QIAGEN) to assess if the contamination was present in the original samples, or introduced later. The V3-V4 region of the 16S rDNA was amplified by PCR using the universal primers 341F/805R^71^, and the Q5 HighFidelity DNA polymerase (NEB). Prior to the addition of the DNA and the primers, the PCR master mix has been treated with 0.02% (v/v) of DNAse I 1 U/mL (Zymo) at room temperature for 15 minutes, followed by DNAse I deactivation at 75°C for 15 minutes, following the manufacturer instruction, to ensure complete decontamination of the master mix ^72^. The PCR has been performed following the manufacturer instructions, using a Tm = 60°C and 30 cycles. Amplicons have been checked on a 1.5% (w/v) agarose gel, and sent for Sanger sequencing with primers 341F and 805R to an external facility. The sequences obtained have been compared with sequences from the GenBank database using BLASTN (NCBI), and EZBioCloud (CJ Bioscience), for the bacterial identification.

### 4.7 Inductively Coupled Plasma Mass Spectrometry (ICP-MS)

One millilitre of the liquid fraction of each sample was treated with nitric acid (final concentration 4% v/v), and the samples were analysed by ICP-MS, to determine concentrations of the elements bioleached from the meteorite. ICP-MS analysis was also carried out on the medium (50% v/v R2A) and fixative (RNAlater).

All samples were analysed for a variety of elements using an Agilent 8900 ICP-MS instrument employing an RF (radio-frequency) forward power of 1550 W, RF matching of 1.8 V, with Argon gas flows of 1.02 L/min and 0.90 L/min for nebuliser and auxiliary flows, respectively. Sample solutions were taken up into a micro mist nebuliser by peristaltic pump at a rate of approximately 1.2 mL/min. Skimmer and sample cones were made of nickel. The instrument was operated in spectrum multi-tune acquisition mode (three replicates runs per sample) for the three isotopes 101Ruthenium, 105Palladium and 195Platinum using Helium mode with a flow rate of He 5 mL/min. To calibrate the instrument, a multi-element calibration standard containing each element was prepared using 1000 mg /L single-element standards (SPE Science, Canada) diluted with 2% (v/v) HNO_3_ (Aristar grade, VWR International, United Kingdom). The limits of detection for each element in He mode were 0.005, 0.005 and 0.007 µg/L for 101Ruthenium, 105Palladium and 195Platinum respectively. Raw ICP-MS data (determined in μg/L) was converted to obtain the absolute quantity of a given element in the culture chamber, taking into account dilution factors applied during ICP-MS analysis.

To determine elemental concentrations in the L-chondrite material, between 25 and 50 mg of homogenised pristine sample (x3) was added to Savillex Teflon vessels. Rock standards (georem standards BCR2, BHVO1 and B-EN) were prepared in the same way. Two blanks were included (i.e., sample without L-chondrite). Three millilitres of double distilled HNO_3_, 2 mL HCl and 0.5 mL HF was added to each of the vessels. HF was added after the other acids to prevent disassociation, formation and precipitation of aluminium fluorides. The HF addition is a necessary step in this protocol, however it compromises the detection of silicon from the rocks, due to its volatilisation. Samples were placed on a hot plate for digestion overnight (temperature of 100– 120 °C) and checked for complete digestion. Samples were evaporated on the hot plate. Five millilitres of 1 M HNO_3_ was added to each vessel. Lids were added and the samples returned to the hot plate for a second digestion step. Samples were further diluted with 2–5% (v/v) HNO_3_ for ICP-MS analysis. Analysis was carried out on a high resolution, sector field, ICP-MS (Nu AttoM). The ICP-MS measurements for elements were performed in low resolution (300), in Deflector jump mode with a dwell time of 1 ms and 3 cycles of 500 sweeps. Data were reported in micrograms of element per gram of chondrite.

### 4.8 Scanning Electron Microscopy and elemental mapping

Representative samples of rock (∼0.3 g) with or without microbial growth were stored in a solution of 3% (v/v) glutaraldehyde in 10 mM HEPES buffer, pH 7.0 for 5 days at 4° C. After this period, stepwise dehydration with graded series of 10, 30, 50, 70, 90, and 100% (v/v) ethanol was performed for 10 min each. Samples were stored at 4° C prior to drying with liquid carbon dioxide in a Polaron E3100 critical point dryer to preserve cell morphologies. Samples were then affixed to SEM Aluminium stubs (Agar Scientific) using a small quantity of conductive carbon glue (Agar Scientific) and coated with 20 nm of gold with a sputter coater (Denton Vacuum) to enhance conductivity for secondary electron imaging.

Further samples were mounted in epoxy resin and polished before carbon coating (Denton BTT-IV carbon evaporation coater) for backscatter electron (BSE) imaging and EDS element mapping. Samples were stored in plastic boxes to prevent dust contamination prior to imaging and analysis using a Carl Zeiss SIGMA HD VP field emission SEM with an Oxford Instruments AZtec EDS system at the School of GeoSciences, University of Edinburgh.

### 4.9 Raman

Raman spectra were recorded with a fibre optic Raman probe and 785 nm stabilized diode laser (Ocean Insight). The probe was mounted to a motorized X-Y-Z translation stage and scanned across the sample surface. Raman spectra were recorded at ca. 0.1 mm lateral resolution and the probe height was adjusted in Z at each point to maximize the Raman signal. The resulting maps were analysed by comparing the Raman peaks at each spectrum to mineral Raman spectra from the RRUFF database to assign a mineral intensity. The broad background fluorescence intensity is the sum of the entire spectrum from 200 – 2000 cm^-1^. The intensity of sharp luminescence peaks is found by summing the spectral region between 1200 – 1600 cm^-1^.

### 4.10 Metabolomics analysis

Polar and non-polar metabolites were analysed using liquid chromatography coupled to high resolution mass spectrometry. Polar metabolites were prepared by diluting the samples a ratio 1:5 (sample/buffer) in extracting buffer (40% v/v MeOH, 40% v/v MeCN and 20% v/v H_2_O) prior to injection. Non-polar metabolites were enriched bz bi-phasic extraction using ethyl-acetate. Metabolites were extracted by vortexing the tubes for 20 min with subsequent spinning down. The organic layer was evaporated and reconstituted in 50% (v/v) MeOH-50% H_2_O (v/v) prior to injection.

During metabolite analysis, a pHILIC column (Merck, Germany) was used to separate polar metabolites, and a Luna C18 (Phenomenex, United States) to separate non-polar metabolites. An Ultimate 3000 HPLC (Thermo Fisher Scientific, Germany) coupled to a Q-Exactive mass spectrometer (Thermo-Fisher Scientific, Germany) operated in polarity switch mode was used. Pooled samples, chemical standards and procedure blanks were also analysed. Detailed description of the methods are included in literature ^73–75^.

Peak detection and integration from Raw data were performed using Compound Discoverer 3.2 (Thermo-Fisher Scientific). An automatic filter set was applied initially to remove features of low quality. Features marked as background signals were removed, with a retention time below one minute, or whose annotated mass diverged by >5ppm from measured mass, were removed. Features with at least two partial matches on reference databases [mzCloud (HighChem LLC), mzVault (Thermo-Fischer Scientific), ChemSpider (Royal Society of Chemistry), and a list of known standards] and full fragmentation data were considered appropriate for further analysis. Partial matches were not discounted as inconsistencies in database entries may affect the match strength without invalidating the annotation. If a feature passed filtering solely based on partial matches, its predicted structure was manually confirmed to be identical to that of the database entry, to ensure correct annotation. Finally, features of insufficient total signal area were removed. Once filtered, all features significantly associated with a condition of interest were identified via differential analysis, with significance determined by p≤0.05. These conditions were: up- or down-regulated in fungal, bacterial, or Consortium culture. Once these lists of hits were produced, the chromatogram of each feature on each list was manually assessed for peak quality. Peaks with a maximum intensity <4x10^6^ counts were removed to ensure sufficient separation from background signals. Features were then assessed individually in greater detail via the metrics produced by Compound Discoverer 3.2. Metrics assayed were both quantitative and qualitative, and consisted of peak areas, group coefficient of variance, adjusted p-values of experimental/control ratios, and the number and identity of positively identified signals per feature. Data analysis and figures were produced using the open source MetaboAnalyst 5.0 program ^76^.

### 4.11 X-Ray Diffraction (XRD)

Pooled samples were analysed using X-ray Diffraction at the School of Geosciences, University of Edinburgh. Fragments of the L-chondrite were gently crushed in a mortar and pestle into a powder. The powder (∼1 g each) was mounted on clean plastic slides. Care was taken to use as little compressional force as possible to minimise preferred mineral grain orientation. The samples were fed into a Bruker D8-Advance X-ray Diffractometer, using a 2-theta configuration in which the X-rays were generated by a Cu-anode X-ray tube operating at 40 kV and a tube current of 40 mA. Diffracted X-rays were detected using a sodium iodide scintillation detector. The samples were scanned from 2 to 60 degrees two theta with a scan rate of 0.02° per second. Resultant diffractograms were compared to the International Centre for Diffraction Data (ICDD) diffractogram database library (2012 issue) using the EVA analysis package. Typically, this procedure gives a detection limit for crystalline phases of approximately 1 wt.%. To quantify mineral abundances in the samples, the diffractograms were subject to Rietveld analysis using the TOPAS software package. This involved identifying the mineral assemblage present by comparing peak positions and heights with those in the powder diffraction database. The TOPAS program then generated a ‘model’ diffraction pattern, calculated from an initial estimated mineral assemblage. The differences between the two are reduced iteratively, which typically takes around 100 iterations, until the model and observed patterns converge, revealing the amounts of the minerals in wt.%.

### 4.12 Data analysis and figure production

Statistical analysis was performed using RStudio 2023.03.0 Build 386 and Microsoft Excel for Microsoft 365 MSO (Version 2303 Build 16.0.16227.20202) 64-bit. Figures were produced using RStudio 2023.03.0 Build 386 and Inkscape 1.1.

## 5. Contribution

RS and CSC conceived and designed the experiment, produced and analysed the experimental data. RS performed and coordinated pre- and post-flight experiments and data analysis, space experiment preparation and wrote the manuscript. RS performed the cell concentration (with partial support of ACW), SEM and elemental mapping analysis. AS produced the RStudio code for the ICP-MS data analysis, supported RS with the sequencing and identification of the contaminants, with the statistical analysis, and with the final draft’s correction. GRB, AG and RS performed the metabolomics analysis. LJE performed the ICP-MS and ICP-OES analysis of the liquid fractions, LP those of the meteorite, GS performed XRD, JH and KRB performed the Raman analysis. NC supported the work with SEM, while MW supported the flow cytometry experiments. RS and CSC performed the pre- and post-flight hardware activities, with the support of SML. MB was the KI scientific referent, AM was the KI project manager and GN was the Bioreactor Express program manager MB, AM and GN prepared and managed the space and Earth bioreactors and containers. All authors approved the manuscript for publication.

## List of nonstandard abbreviations

ISS: International Space Station, under microgravity condition
Earth: Ground controls, under terrestrial gravity condition
µg: Microgravity
PGEs: Platinum group elements
REEs: Rare earth elements
ISRU: *In situ* resource utilisation
BMR: BioMining Reactor
EC: Experiment Container
EU: Experiment Unit
%_NB_: Element amount as percentage of sample / non-biological sample
%_M_: Element amount as percentage of sample / total in the meteorite rock

## 6. Acknowledgements and funding

RS and CSC were supported by United Kingdom Science and Technology Facilities Council under grant ST/V000586/1. RS was partially supported by Leverhulme Trust under grant ECF-2021-185. Edinburgh-Rice Strategic Collaboration Awards supported JH, CSC, RS and KRB with the Raman analysis. We thank SkyFall Meteorites for the L-chondrite sample. We are thankful to Stefano Pellari, Ramon Nartallo, David Zolesi and Alessandro Donati (Kayser Italia Srl, and Kayser Space Ltd) for their support, to Robert Bunker (Meritics Ltd) for the meteorite surface area analysis, and to Stephen Mitchell (University of Edinburgh) for the support on CPD procedures. Flow cytometry data were generated within the Flow Cytometry and Cell Sorting Facility in Ashworth, King’s Buildings (University of Edinburgh), supported by funding from Wellcome and the University of Edinburgh. We thank NASA astronaut Michael Scott Hopkins for taking care of BioAsteroid onboard the ISS, Nic Odling (University of Edinburgh) for the support with the XRD, Rebecca D. Prescott (University of Mississippi, University of Hawai‘i at Mānoa, NASA JSC) for helping with the fixative choice, Virginia Echavarri-Bravo (University of Edinburgh) for useful discussions on flow cytometry, Diana, Sean (and Violet) Marosi-McMahon for their support on the statistical analysis.

## 8. References

1. Verseux, C. et al. Sustainable life support on Mars - The potential roles of cyanobacteria. Int. J. Astrobiol. 15, 65–92 (2016).

2. Gumulya, Y., Zea, L. & Kaksonen, A. H. *In situ* resource utilisation: The potential for space biomining. Miner. Eng. 176, 107288 (2022).

3. Averesch, N. J. H. Choice of Microbial System for *In-Situ* Resource Utilization on Mars. Front. Astron. Sp. Sci. 8, 1–7 (2021).

4. Santomartino, R. et al. Toward sustainable space exploration: a roadmap for harnessing the power of microorganisms. Nat. Commun. 14, 1–11 (2023).

5. Cockell, C. S. Geomicrobiology beyond Earth: Microbe-mineral interactions in space exploration and settlement. Trends Microbiol. 18, 308–314 (2010).

6. Santomartino, R., Zea, L. & Cockell, C. S. The smallest space miners: principles of space biomining. Extremophiles 26, 1–19 (2022).

7. Housen, K. R., Wilkening, L. L., Chapman, C. R. & Greenberg, R. Asteroidal regoliths. Icarus 39, 317–351 (1979).

8. Rubin, A. E. Mineralogy of meteorite groups. Meteorit. Planet. Sci. 32, 231–247 (1997).

9. Ross, S. D. Near-Earth Asteroid Mining. in Space Industry Report 1–24 (2001).

10. Klas, M. et al. Biomining and methanogenesis for resource extraction from asteroids. Space Policy 34, 18–22 (2015).

11. Steenstra, E. S. et al. An experimental assessment of the potential of sulfide saturation of the source regions of eucrites and angrites: Implications for asteroidal models of core formation, late accretion and volatile element depletions. Geochim. Cosmochim. Acta 269, 39–62 (2020).

12. Kettler, P. B. Platinum group metals in catalysis: Fabrication of catalysts and catalyst precursors. Org. Process Res. Dev. 7, 342–354 (2003).

13. Hein, A. M., Matheson, R. & Fries, D. A techno-economic analysis of asteroid mining. Acta Astronaut. 168, 104–115 (2020).

14. Gertsch, R. E. Asteroid mining. in Space resources: materials 111–120 (1992).

15. Cockell, C. S. Synthetic geomicrobiology: Engineering microbe-mineral interactions for space exploration and settlement. Int. J. Astrobiol. 10, 315–324 (2011).

16. Averesch, N. J. H. et al. Microbial biomanufacturing for space-exploration—what to take and when to make. Nat. Commun. 14, 1–10 (2023).

17. Nangle, S. N. et al. The case for biotech on Mars. Nat. Biotechnol. 38, 401–407 (2020).

18. Berliner, A. J. et al. Space bioprocess engineering on the horizon. Commun. Eng. 1, 13 (2022).

19. Schippers, A. et al. Biomining: Metal Recovery from Ores with Microorganisms. Geobiotechnology I. Adv. Biochem. Eng. 141, 1–47 (2013).

20. Roberto, F. F. & Schippers, A. Progress in bioleaching: part B, applications of microbial processes by the minerals industries. Appl. Microbiol. Biotechnol. 106, 5913–5928 (2022).

21. Johnson, D. B., Grail, B. M. & Hallberg, K. B. A new direction for biomining: Extraction of metals by reductive dissolution of oxidized ores. Minerals 3, 49–58 (2013).

22. Brune, K. D. & Bayer, T. S. Engineering microbial consortia to enhance biomining and bioremediation. Front. Microbiol. 3, 1–6 (2012).

23. Brandl, H., Bosshard, R. & Wegmann, M. Computer-munching microbes: Metal leaching from electronic scrap by bacteria and fungi. Process Metall. 9, 569–576 (1999).

24. Rohwerder, T., Gehrke, T., Kinzler, K. & Sand, W. Bioleaching review part A: Progress in bioleaching: Fundamentals and mechanisms of bacterial metal sulfide oxidation. Appl. Microbiol. Biotechnol. 63, 239–248 (2003).

25. Noël, N., Florian, B. & Sand, W. AFM & EFM study on attachment of acidophilic leaching organisms. Hydrometallurgy 104, 370–375 (2010).

26. Adeleke, R., Cloete, E. & Khasa, D. Isolation and identification of iron ore-solubilising fungus. S. Afr. J. Sci. 106, 1–6 (2010).

27. Din, G. et al. Characterization of Organic Acid Producing *Aspergillus tubingensis* FMS1 and its Role in Metals Leaching from Soil. Geomicrobiol. J. 37, 336–344 (2020).

28. Barnett, M. J., Palumbo-Roe, B. & Gregory, S. P. Comparison of heterotrophic bioleaching and ammonium sulfate ion exchange leaching of rare earth elements from a Madagascan ion-adsorption clay. Minerals 8, 1–11 (2018).

29. Gadd, G. M. Fungal production of citric and oxalic acid: Importance in metal speciation, physiology and biogeochemical processes. Advances in Microbial Physiology vol. 41 (Elsevier Masson SAS, 1999).

30. Cockell, C. S. & Santomartino, R. Mining and Microbiology for the Solar System Silicate and Basalt Economy. in *In Space Manufacturing Resources: Earth and Planetary Exploration Applications* (eds. Hessel, V., Stoudemire, J., Miyamoto, H. & Fisk, I. D.) 163–185 (Wiley, 2022). 10.1002/9783527830909.ch8.

31. Cockell, C. S. et al. Microbially-Enhanced Vanadium Mining and Bioremediation Under Micro- and Mars Gravity on the International Space Station. Front. Microbiol. 12, 663 (2021).

32. Cockell, C. S. et al. Space station biomining experiment demonstrates rare earth element extraction in microgravity and Mars gravity. Nat. Commun. 11, 1–12 (2020).

33. Byloos, B. et al. The impact of space flight on survival and interaction of *Cupriavidus metallidurans* CH34 with basalt, a volcanic moon analog rock. Front. Microbiol. 8, 1–14 (2017).

34. Castelein, S. M. et al. Iron can be microbially extracted from Lunar and Martian regolith simulants and 3D printed into tough structural materials. PLoS One 16, 1–21 (2021).

35. Figueira, J. et al. Biomining of Lunar regolith simulant EAC-1A with the fungus *Penicillium simplicissimum*. Res. Sq. (2023) 10.21203/rs.3.rs-2909117/v1.

36. Waajen, A. C., Prescott, R. & Cockell, C. S. Meteorites as Food Source on Early Earth: Growth, Selection, and Inhibition of a Microbial Community on a Carbonaceous Chondrite. Astrobiology 22, (2022).

37. Milojevic, T. et al. Exploring the microbial biotransformation of extraterrestrial material on nanometer scale. Sci. Rep. 9, 1–11 (2019).

38. Milojevic, T. et al. Chemolithotrophy on the Noachian Martian breccia NWA 7034 via experimental microbial biotransformation. Commun. Earth Environ. 2, (2021).

39. Reddy, G. S. N. & Garcia-Pichel, F. *Sphingomonas mucosissima* sp. nov. and *Sphingomonas desiccabilis* sp. nov., from biological soil crusts in the Colorado Plateau, USA. Int. J. Syst. Evol. Microbiol. 57, 1028–1034 (2007).

40. Asaf, S., et al., *Sphingomonas*: from diversity and genomics to functional role in environmental remediation and plant growth. Critical Reviews in Biotechnology, 40:2, 138–152, (2020).

41. Ambreen, N., Bhatti, H. N. & Bhatti, T. M. Bioleaching of Bauxite by *Penicillium simplicissimum*. J. Biol. Sci. 2, 793–796 (2002).

42. Franz, A., Burgstaller, W. & Schinner, F. Leaching with *Penicillium simplicissimum*: Influence of metals and buffers on proton extrusion and citric acid production. Appl. Environ. Microbiol. 57, 769–774 (1991).

43. Amiri, F., Yaghmaei, S. & Mousavi, S. M. Bioleaching of tungsten-rich spent hydrocracking catalyst using *Penicillium simplicissimum*. Bioresour. Technol. 102, 1567– 1573 (2011).

44. Rawlings, D. E. & Johnson, D. B. The microbiology of biomining: Development and optimization of mineral-oxidizing microbial consortia. Microbiology 153, 315–324 (2007).

45. McCoy-West, A. J., Millet, M. A. & Burton, K. W. The neodymium stable isotope composition of the silicate Earth and chondrites. Earth Planet. Sci. Lett. 480, 121–132 (2017).

46. Gaft, M., Reisfeld, R. & Panczer, G. Modern Luminescence Spectroscopy of Minerals and Materials. (Springer Mineralogy, 2015). doi:10.1007/978-3-319-24765-6.

47. Petrikkou, E. et al. Inoculum standardization for antifungal susceptibility testing of filamentous fungi pathogenic for humans. J. Clin. Microbiol. 39, 1345–1347 (2001).

48. Bellenberg, S. et al. Biofilm formation, communication and interactions of leaching bacteria during colonization of pyrite and sulfur surfaces. Res. Microbiol. 165, 773–781 (2014).

49. Zuo, R., Kus, E., Mansfeld, F. & Wood, T. K. The importance of live biofilms in corrosion protection. Corros. Sci. 47, 279–287 (2005).

50. Jayaraman, A., Sun, A. K. & Wood, T. K. Characterization of axenic *Pseudomonas fragi* and *Escherichia coli* biofilms that inhibit corrosion of SAE 1018 steel. J. Appl. Microbiol. 84, 485–492 (1998).

51. Zuo, R. Biofilms: Strategies for metal corrosion inhibition employing microorganisms. Appl. Microbiol. Biotechnol. 76, 1245–1253 (2007).

52. Stevens, A. H. et al. Growth, Viability, and Death of Planktonic and Biofilm *Sphingomonas desiccabilis* in Simulated Martian Brines. Astrobiology 19, 87–98 (2019).

53. Reed, D. W., Fujita, Y., Daubaras, D. L., Jiao, Y. & Thompson, V. S. Bioleaching of rare earth elements from waste phosphors and cracking catalysts. Hydrometallurgy 166, 34–40 (2016).

54. Vailati, A. et al. Diffusion in liquid mixtures. npj Microgravity 9, 1–8 (2023).

55. Huang, B., Li, D. G., Huang, Y. & Liu, C. T. Effects of spaceflight and simulated microgravity on microbial growth and secondary metabolism. Mil. Med. Res. 5, 1–14 (2018).

56. De Gelder, J. et al. Raman spectroscopic analysis of *Cupriavidus metallidurans* LMG 1195 (CH34) cultured in low-shear microgravity conditions. Microgravity Sci. Technol. 21, 217–223 (2009).

57. Saleh, D. K., Abdollahi, H., Noaparast, M., Nosratabad, A. F. & Tuovinen, O. H. Dissolution of Al from metakaolin with carboxylic acids produced by *Aspergillus niger*, *Penicillium bilaji*, *Pseudomonas putida*, and *Pseudomonas koreensis*. Hydrometallurgy 186, 235–243 (2019).

58. Panhwar, Q. A., Naher, U. A., Shamshuddin, J., Othman, R. & Latif, M. A. Biochemical and molecular characterization of potential phosphate-solubilizing bacteria in acid sulfate soils and their beneficial effects on rice growth. PLoS One 9, (2014).

59. Shang, R. Palladium-Catalyzed Decarboxylative Coupling of Potassium Oxalate Monoester with Aryl and Alkenyl Halides. in New Carbon–Carbon Coupling Reactions Based on Decarboxylation and Iron-Catalyzed C–H Activation 49–61 (Springer Nature Singapore Pte Ltd., 2017). doi:10.1007/978-981-10-3193-9_2.

60. Li, X. & Binnemans, K. Oxidative Dissolution of Metals in Organic Solvents. Chem. Rev. 121, 4506–4530 (2021).

61. Li, C. et al. Assessment of 2,2,6,6-tetramethyl-4-piperidinol-based amine N-halamine-labeled silica nanoparticles as potent antibiotics for deactivating bacteria. Colloids Surfaces B Biointerfaces 126, 106–114 (2015).

62. Furumura, S. et al. Identification and Functional Characterization of Fungal Chalcone Synthase and Chalcone Isomerase. J. Nat. Prod. 86, 398–405 (2023).

63. Oenbrink, G. & Schiffer, T. Cyclododecanol, Cyclododecanone, and Laurolactam. Ullmann’s Encycl. Ind. Chem. 1–5 (2009) doi:10.1002/14356007.a08_201.pub2.

64. Santomartino, R. et al. No Effect of Microgravity and Simulated Mars Gravity on Final Bacterial Cell Concentrations on the International Space Station: Applications to Space Bioproduction. Front. Microbiol. 11, 1–15 (2020).

65. Volger, R. et al. Mining moon & mars with microbes: Biological approaches to extract iron from Lunar and Martian regolith. Planet. Space Sci. 184, 104850 (2020).

66. Prescott, R. D. et al. Including Descriptions of Three Novel Bacterial Species Isolated from Mars Analog Sites of Cultural Relevance. Astrobiology 23, 1–20 (2023).

67. Rasoulnia, P., Mousavi, S. M., Rastegar, S. O. & Azargoshasb, H. Fungal leaching of valuable metals from a power plant residual ash using *Penicillium simplicissimum*: Evaluation of thermal pretreatment and different bioleaching methods. Waste Manag. 52, 309–317 (2016).

68. Welten, K. et al. The L3-6 chondritic regolith breccia Northwest Africa (NWA) 869: (II) Noble gases and cosmogenic radionuclides. Meteorit. Planet. Sci. 46, 970–988 (2011).

69. Metzler, K. et al. The L3-6 chondritic regolith breccia Northwest Africa (NWA) 869: (I) Petrology, chemistry, oxygen isotopes, and Ar-Ar age determinations. Meteorit. Planet. Sci. 46, 652–680 (2011).

70. Loudon, C. M. et al. BioRock: new experiments and hardware to investigate microbe– mineral interactions in space. Int. J. Astrobiol. 17, 303–313 (2018).

71. Herlemann, D. P. R. et al. Transitions in bacterial communities along the 2000 km salinity gradient of the Baltic Sea. ISME J. 5, 1571–1579 (2011).

72. Heininger, A. et al. DNase pretreatment of master mix reagents improves the validity of universal 16S rRNA gene PCR results. J. Clin. Microbiol. 41, 1763–1765 (2003).

73. Chetwynd, A. J., Dunn, W. B. & Rodriguez-Blanco, G. Metabolomics: From Fundamentals to Clinical Applications. in Metabolomics: From Fundamentals to Clinical Applications, Advances in Experimental Medicine and Biology (ed. Sussulini, A.) vol. 965 19–44 (Springer International Publishing AG 2017, 2017).

74. MacKay, G. M., Zheng, L., Van Den Broek, N. J. F. & Gottlieb, E. Analysis of Cell Metabolism Using LC-MS and Isotope Tracers. in Methods in Enzymology vol. 561 171– 196 (Elsevier Inc., 2015).

75. Sarafian, M. H. et al. Objective set of criteria for optimization of sample preparation procedures for ultra-high throughput untargeted blood plasma lipid profiling by ultra performance liquid chromatography-mass spectrometry. Anal. Chem. 86, 5766–5774 (2014).

76. Pang, Z. et al. MetaboAnalyst 5.0: Narrowing the gap between raw spectra and functional insights. Nucleic Acids Res. 49, W388–W396 (2021).

