## Supplementary information for "Testing microbial biomining from asteroidal material onboard the International Space Station"

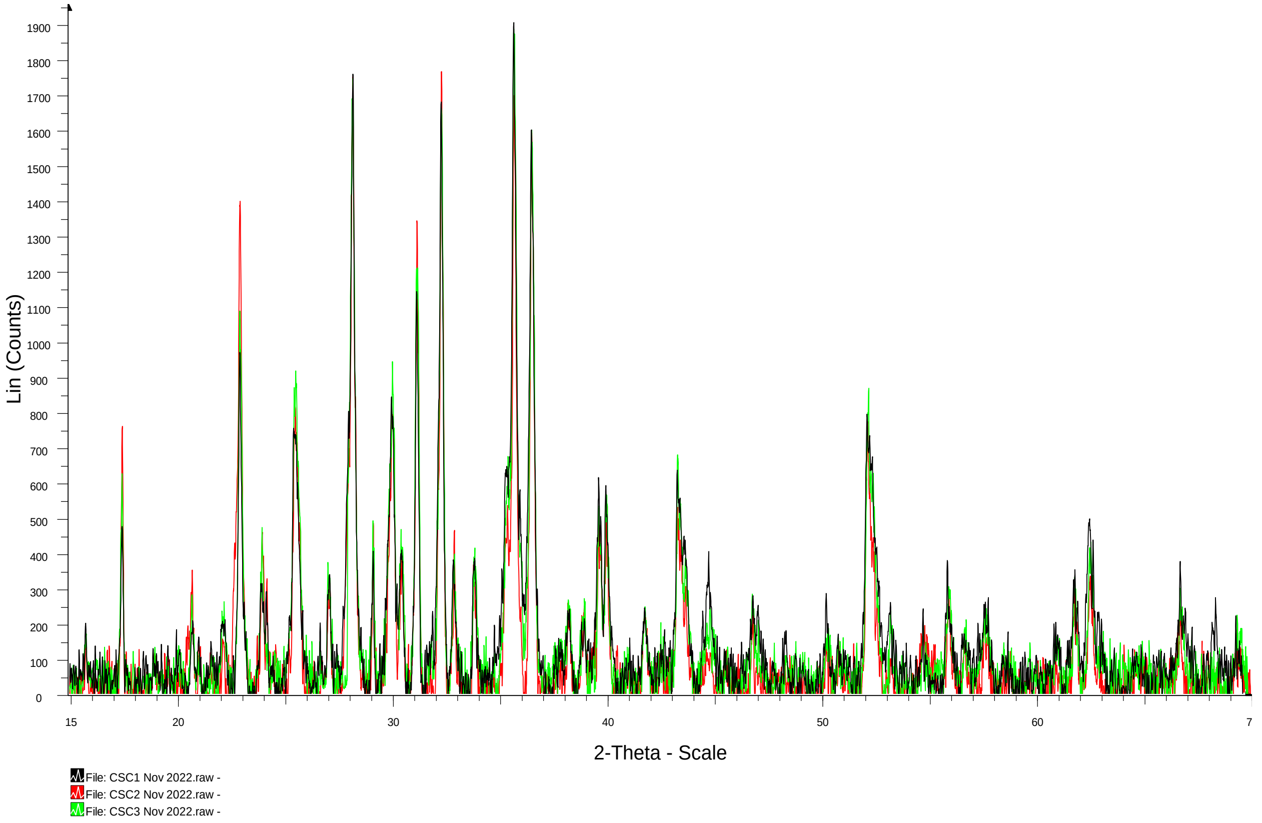


***S1*** XRD analysis of the meteorite rock (L-chondrite) used in this experiment. Comparison between 3 replicates (black, red and green) are shown here.

| Sample | OD$_{600}$ | | OD$_{530}$ | | Flow cytometry (Cell/mL) | | | |
|---|---|---|---|---|---|---|---|---|
| | | | | | Baclight Green (bacterial dye) | | Calcofluor White (fungal dye) | |
| | ISS | Earth | ISS | Earth | ISS | Earth | ISS | Earth |
| Non-biological | 0.15±0.01 | 0.28±0.05 | 0.21±0.01 | 0.47±0.15 | (6.66±0.37)x10$^{7}$ | (7.77±4.75)x10$^{7}$ | (6.33±1.19)x10$^{4}$ | (2.68±1.94)x10$^{4}$ |
| ***S. desiccabilis*** | 0.21±0.08 | 0.64±0.05 | 0.35±0.10 | 1.18±0.15 | (3.97±0.06)x10$^{9}$ | (3.27±0.32)x10$^{9}$ | (3.15±0.75)x10$^{6}$ | (2.80±0.70)x10$^{6}$ |
| ***P. simplicissimum*** | 0.03±0.00 | 0.29±0.12 | 0.06±0.01 | 0.44±0.14 | (2.80±0.98) x10$^{8}$ | (3.07±1.22)x10$^{8}$ | (3.50±1.99)x10$^{6}$ | (6.37±2.11)x10$^{7}$ |
| Consortium | 0.04±0.00 | 0.46±0.02 | 0.06±0.01 | 0.74±0.06 | (6.47±1.98)x10$^{8}$ | (3.64±0.14)x10$^{9}$ | (1.43±0.34)x10$^{6}$ | (7.69±1.33)x10$^{6}$ |
| Diluent (PBS) | | | | | - | (4.90±2.40)x10$^{4}$ | - | (4.50±2.50)x10$^{2}$ |

***S2*** Spectrometric analysis (optical density, λ = 600nm, 530 nm) and flow cytometry analysis measured from the liquid fraction of the space (ISS) and ground control (Earth) samples of the BioAsteroid samples. A wavelength of 530 nm, instead of the classical 600 nm, was used as an additional measure for measuring filamentous fungi growth (Petrikkou et al., 2001). For flow cytometry, values represent the average events counted in 100 µL of sample stained with Baclight green (specific for bacterial cells) and calcofluor white (specific for the fungal cells). Values are indicated as mean±standard error.

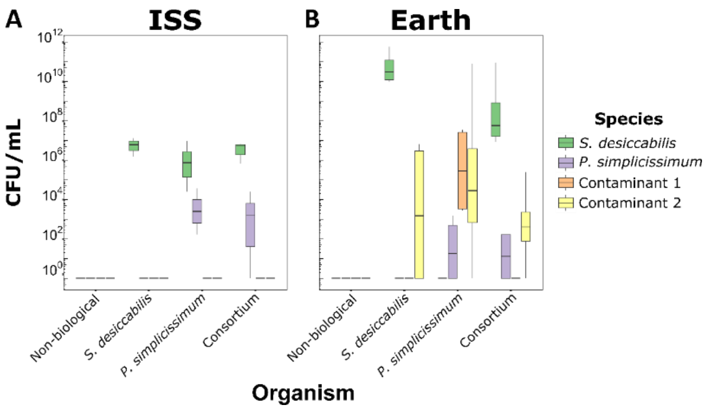


***S3*** Final cell concentrations measured in the liquid fraction of the space (ISS, A) and ground control (Earth, B) samples in terms of CFU/mL. CFU concentration of *S desiccabilis*, *P. simplicissimum*, Contaminant 1 and Contaminant 2 species are indicated.

| | pH | |
|---|---|---|
| **Organism** | **ISS** | **Earth** |
| **Non-biological** | 7.21±0.03 | 7.35±0.02 |
| ***S. desiccabilis*** | 7.34±0.01 | 7.38±0.01 |
| ***P. simplicissimum*** | 7.33±0.03 | 7.35±0.02 |
| **Consortium** | 7.43±0.01 | 7.43±0.01 |
| **R2A 50%** | - | 7.2 |
| **RNAlater** | - | 4.87±0.01 |
| **R2A 50% + RNAlater (1:5 v/v)** | - | 5.16±0.04 |

***S4*** pH of the liquid fraction of the space (ISS) and ground control (Earth) samples. Values are indicated as mean±standard error. As pH is one of the potential influencing factors in leaching, pH of the liquid fractions was measured after spaceflight. The normal Q-Q plot (data not shown) and the Shapiro-Wilks test suggested the normality of the pH data (W = 0.94836, p-value = 0.1528), hence the parametric ANOVA test was performed to compare pH values. The variables *organism* (F = 15.170, p-value = 1.45 x $10^{-5}$), the *gravity* (F = 11.945, p-value = 0.002), as well as the interaction between the two (F = 5.044, p-value = 0.008) had an effect on the pH values. Pairwise comparison (post-hoc Tukey test) between non-biological and biological samples on the ISS had a p-value < 0.05 in all cases (Non-bio/*S. desiccabilis*: p-value = 0.012; Non-bio/*P. simplicissimum*: p-value = 0.026; Non-bio/Consortium: p-value = $9.36 \times 10^{-6}$), while similar comparisons on Earth reported p-values > 0.05. Comparison between non-biological samples on the ISS and on Earth had a p-value = 0.001. Comparison between *P. simplicissimum* and Consortium on the ISS had a p-value = 0.039. All other comparisons had a p-value > 0.05. The rock may sequester protons and buffer the solution, and this effect might be different under space and Earth conditions. This suggests that pH does not have a bioleaching role for *S. desiccabilis* and *P. simplicissimum*. Similar observations were made for rare earth element and vanadium extraction in space by *S. desiccabilis*, where bioleaching occurred under circumneutral pH (Cockell et al., 2020, 2021). However, the pH values reported were likely buffered by the presence of the fixative (RNAlater), thus impairing our capacity to drive definitive conclusion on the effects of pH in our system.

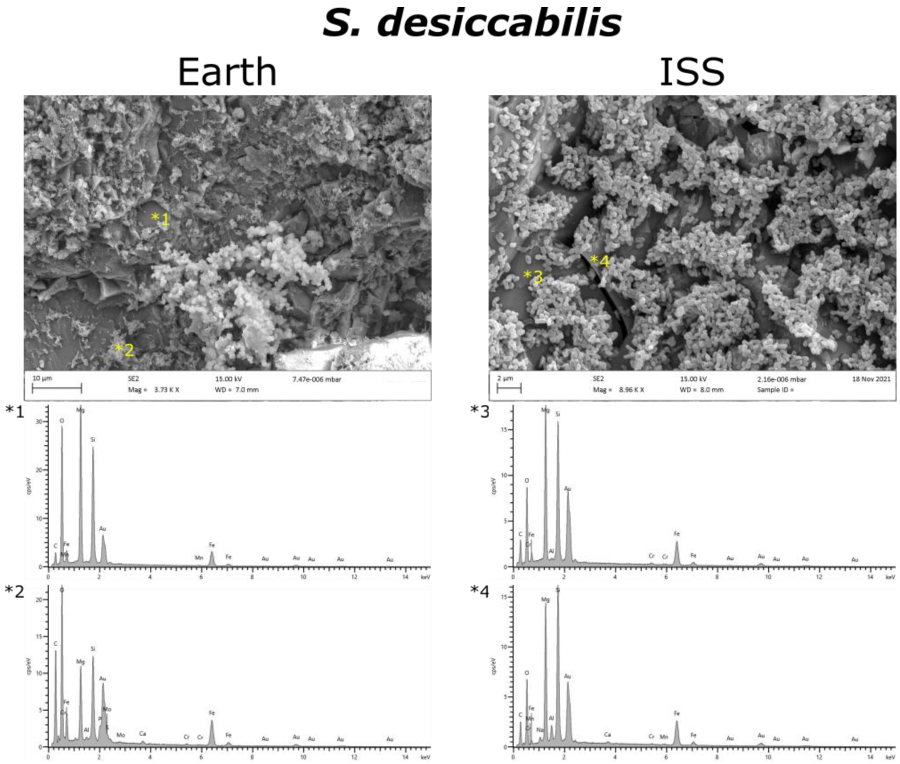


***S5*** Scanning electron microscopy images of *S. desiccabilis* biofilm formed on top of the meteorite rock fragments for ground (Earth) and space (ISS) samples. Asterisks indicate the points were energy-dispersive X-ray spectroscopy (EDS) spectra were measured. Gold peaks are from the gold sample coating.

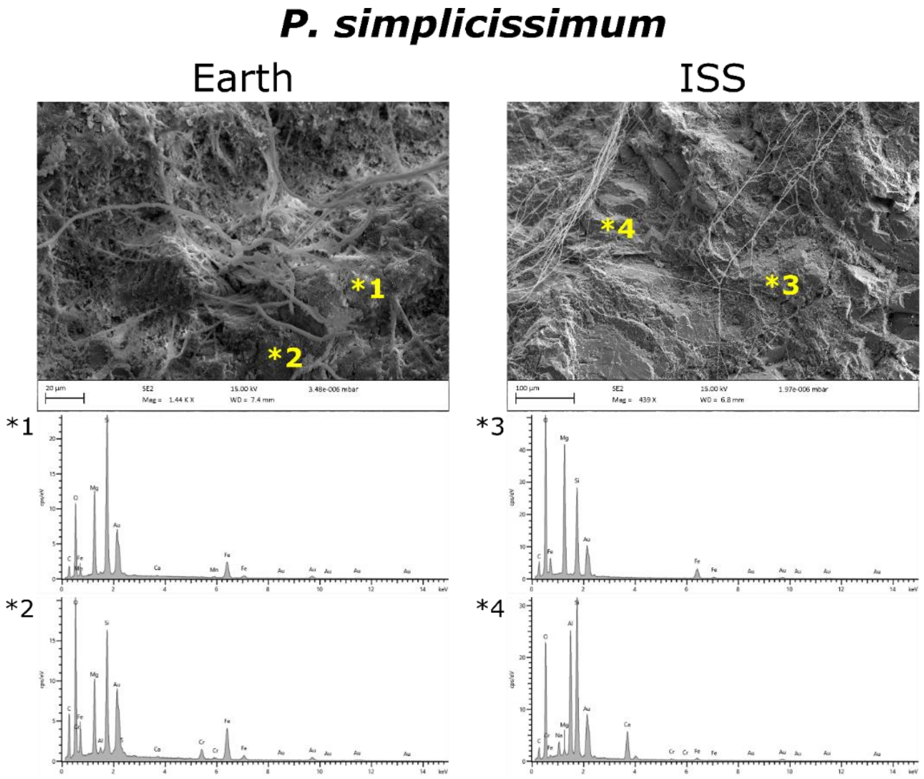


***S6*** Scanning electron microscopy images of *P. simplicissimum* mycelium formed on top of the meteorite rock fragments for ground (Earth) and space (ISS) samples. Asterisks indicate the points were EDS spectra were measured. Gold peaks are from the gold sample coating.

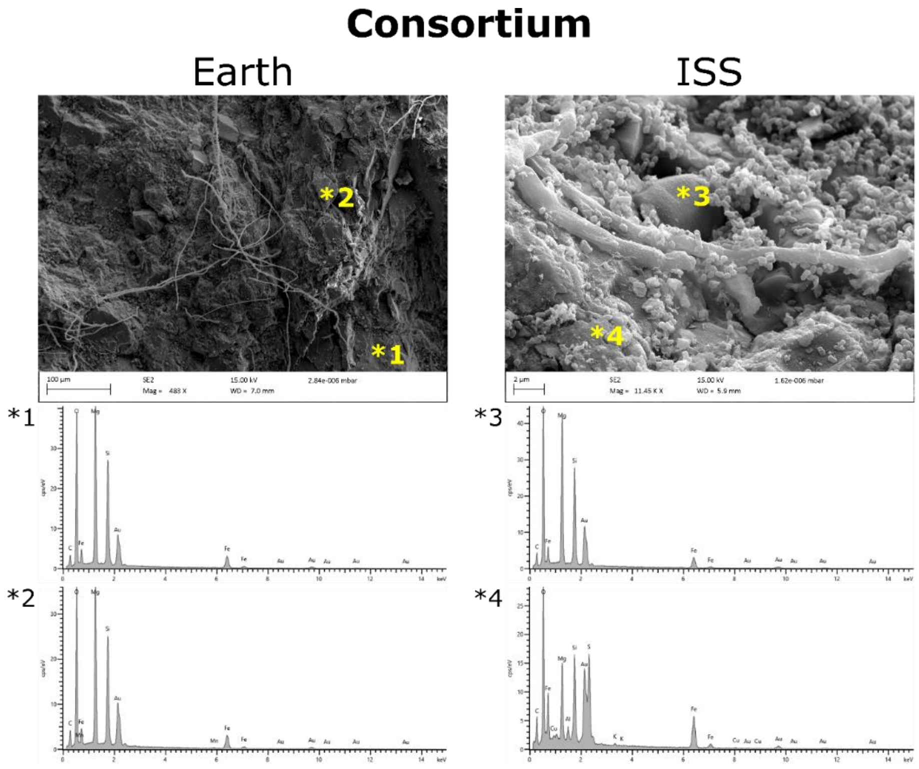


***S7*** Scanning electron microscopy images of bacterial biofilm and fungal mycelium formation on meteorite rock fragments for ground (Earth) and space (ISS) consortium samples. Asterisks indicate the points were EDS spectra were measured. Gold peaks are from the gold sample coating.

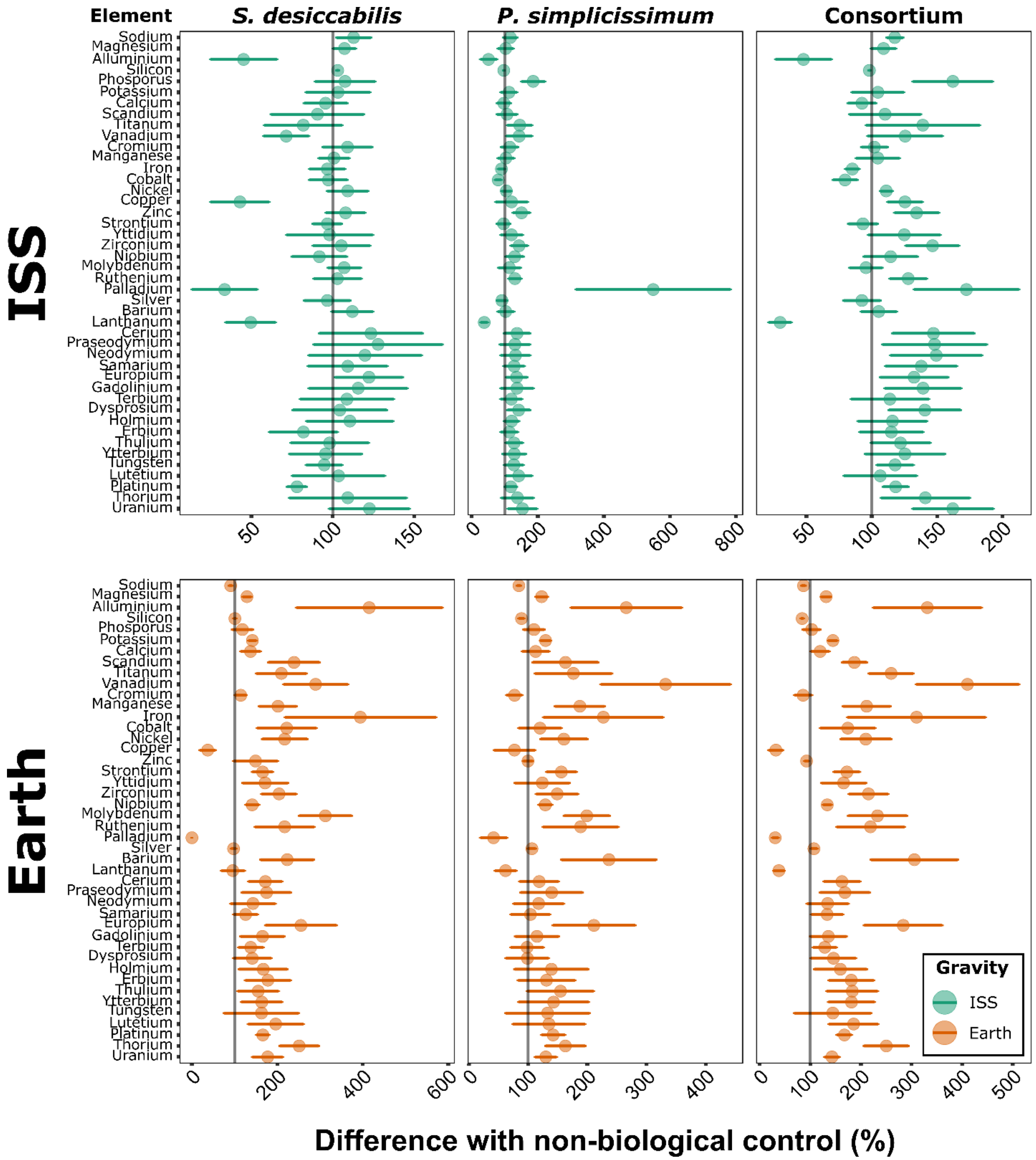


***S8* Analysis of biomining for 44 elements on the ISS and on Earth.** For each element, represented in order of atomic number, bioleaching data are shown as percentage differences with the non-biological control. Upper panels indicate data from the ISS samples, lower panels from the Earth samples. Circles indicates mean values; error bars indicate standard error; vertical grey lines indicate 100% values (i.e., same bioleaching efficiency as non-biological samples).

| | | *Organism* | | *Gravity* | | *Organism x Gravity* | |
|---|---|---|---|---|---|---|---|
| **#** | **Element** | **F** | **p value** | **F** | **p value** | **F** | **p value** |
| *1* | Sodium | 0.830 | 0.492 | 15.282 | **0.001** | 3.384 | **0.036** |
| *2* | Aluminium | 0.254 | 0.858 | 4.730 | **0.041** | 5.054 | **0.008** |
| *3* | Silicon | 10.449 | **0.0002** | 18.389 | **0.0003** | 3.172 | **0.044** |
| *4* | Phosphorus | 2.467 | 0.089 | 0.166 | 0.688 | 3.344 | **0.038** |
| *5* | Potassium | 5.778 | **0.005** | 7.354 | **0.013** | 2.720 | 0.069 |
| *6* | Scandium | 2.028 | 0.139 | 29.871 | **0.00002** | 1.354 | 0.283 |
| *7* | Vanadium | 10.941 | **0.0001** | 3.548 | 0.073 | 3.488 | **0.033** |
| *8* | Manganese | 6.042 | **0.004** | 1.18910 | 0.287 | 2.745 | 0.067 |
| *9* | Iron | 9.908 | **0.00025** | 29.340 | **0.00002** | 6.908 | **0.002** |
| *10* | Cobalt | 4.731 | **0.011** | 15.279 | **0.001** | 3.138 | **0.046** |
| *11* | Nickel | 8.519 | **0.001** | 29.196 | **0.00002** | 2.060 | 0.135 |
| *12* | Copper | 1.536 | 0.233 | 6,473 | **0.019** | 0.695 | 0.565 |
| *13* | Strontium | 4.118 | **0.018** | 7.106 | **0.014** | 3.057 | **0.050** |
| *14* | Zirconium | 11.172 | **0.0001** | 5.442 | **0.029** | 2.187 | 0.118 |
| *15* | Molybdenum | 5.745 | **0.005** | 10.495 | **0.004** | 2.668 | 0.073 |
| *16* | Ruthenium | 6.744 | **0.002** | 15.859 | **0.001** | 1.506 | 0.241 |
| *17* | Palladium | 12.148 | **0.0001** | 10.405 | **0.004** | 6.709 | **0.002** |
| *18* | Barium | 6.041 | **0.004** | 0.167 | 0.687 | 2.862 | **0.060** |
| *19* | Europium | 6.051 | **0.004** | 12.202 | **0.002** | 0.764 | 0.526 |
| *20* | Erbium | 2.010 | 0.142 | 21.970 | **0.0001** | 1.530 | 0.235 |
| *21* | Lutetium | 3.783 | **0.025** | 23.987 | **0.0001** | 1.690 | 0.198 |
| *22* | Platinum | 9.992 | **0.00024** | 30.137 | **0.00002** | 7.478 | **0.001** |

***S9*** Statistical values calculated with ANOVA for the raw concentrations of the 22 elements whose p-value was ≤ 0.05 for at least one of the variables analysed (*Organism*, *Gravity* or their interaction), and for at least one biologically relevant pairwise comparison. In pale yellow the PGEs analysed in this work; in bold p-values ≤ 0.05.

| | | | ISS | | | | Earth | | | |
|---|---|---|---|---|---|---|---|---|---|---|
| | | | NB | SD | PS | CN | NB | SD | PS | CN |
| Ruthenium | ISS | NB | | | | | | | | |
| | | SD | 1 | | | | | | | |
| | | PS | 0.76 | 0.83 | | | | | | |
| | | CN | 0.83 | 0.89 | 1 | | | | | |
| | Earth | NB | 0.54 | 0.04 | 0.0009 | 0.0012 | | | | |
| | | SD | 1 | 1 | 0.59 | 0.68 | 0.042 | | | |
| | | PS | 0.96 | 0.94 | 0.22 | 0.28 | 0.212 | 0.99 | | |
| | | CN | 1 | 1 | 0.62 | 0.71 | 0.038 | 1 | 0.99 | |
| Palladium | ISS | NB | | | | | | | | |
| | | SD | 0.99 | | | | | | | |
| | | PS | 0.73 | 0.58 | | | | | | |
| | | CN | 0.99 | 0.99 | 0.86 | | | | | |
| | Earth | NB | 0.000624 | 0.000327 | 0.05 | 0.001264 | | | | |
| | | SD | 0.99 | 1 | 0.44 | 0.99 | 7.02 x $10^{-5}$ | | | |
| | | PS | 0.62 | 0.47 | 1 | 0.79 | 0.03 | 0.31 | | |
| | | CN | 0.92 | 0.81 | 0.99 | 0.98 | 0.006 | 0.67 | 0.99 | |
| Platinum | ISS | NB | | | | | | | | |
| | | SD | 0.44 | | | | | | | |
| | | PS | 0.65 | 0.016 | | | | | | |
| | | CN | 0.66 | 0.016 | 1.00 | | | | | |
| | Earth | NB | 0.001 | 0.20 | 0.00001 | 0.00001 | | | | |
| | | SD | 0.97 | 0.91 | 0.13 | 0.13 | 0.005 | | | |
| | | PS | 0.32 | 1.00 | 0.007 | 0.01 | 0.15 | 0.83 | | |
| | | CN | 0.98 | 0.88 | 0.14 | 0.15 | 0.004 | 1.00 | 0.79 | |

***S10*** P-values from the pairwise comparison of the raw concentrations with Tukey post-hoc test for the PGEs (ruthenium, palladium and platinum). Green shadowed cells indicate p-values < 0.05 from biologically relevant comparison. Red text indicates the comparisons that could have been influenced by the anomalies reported in S3. NB = non-biological; SD = *S. desiccabilis*; PS = *P. simplicissimum*; CN = Consortium.

| | Ruthenium | | | | | | Palladium | | | | | | Platinum | | | | | |
|---|---|---|---|---|---|---|---|---|---|---|---|---|---|---|---|---|---|---|
| | ng/mL (ppb) | | $\%_{M}$ | | $\%_{NB}$ | | ng/mL (ppb) | | $\%_{M}$ | | $\%_{NB}$ | | ng/mL (ppb) | | $\%_{M}$ | | $\%_{NB}$ | |
| | ISS | Earth | ISS | Earth | ISS | Earth | ISS | Earth | ISS | Earth | ISS | Earth | ISS | Earth | ISS | Earth | ISS | Earth |
| ***NB*** | 0.22 ±0.02 | 0.10 ±0.03 | 14.78 ±2.19 | 6.59 ±2.13 | 100.00 ±13.23 | 100.00 ±42.73 | 0.05 ±0.01 | 0.69 ±0.10 | 2.17 ±0.58 | 29.47 ±6.47 | 100.00 ±29.82 | 100.00 ±20.65 | 0.22 ±0.02 | 0.12 ±0.01 | 0.24 ±0.04 | 0.13 ±0.02 | 100.00 ±10.07 | 100.00 ±12.25 |
| ***SD*** | 0.23 ±0.02 | 0.21 ±0.02 | 15.20 ±2.40 | 14.32 ±2.30 | 102.87 ±14.67 | 217.12 ±69.93 | 0.02 ±0.01 | 0.00 ±0.00 | 0.73 ±0.43 | 0.05 ±0.05 | 33.55 ±20.11 | 0.18 ±0.18 | 0.17 ±0.00 | 0.20 ±0.01 | 0.19 ±0.02 | 0.22 ±0.03 | 78.03 ±5.91 | 166.09 ±12.25 |
| ***PS*** | 0.29 ±0.03 | 0.19 ±0.03 | 19.29 ±3.22 | 12.44 ±2.37 | 130.56 ±19.94 | 188.64 ±63.81 | 0.29 ±0.10 | 0.29 ±12.3 | 11.91 ±4.83 | 12.25 ±6.70 | 549.34 ±234.43 | 41.58 ±22.53 | 0.26 ±0.03 | 0.17 ±0.02 | 0.29 ±0.05 | 0.19 ±0.03 | 118.39 ±17.63 | 142.16 ±20.14 |
| ***CN*** | 0.28 ±0.02 | 0.22 ±0.01 | 18.89 ±2.46 | 14.44 ±1.88 | 127.84 ±14.24 | 218.90 ±67.45 | 0.09 ±0.01 | 0.21 ±0.03 | 3.74 ±0.72 | 9.03 ±1.91 | 172.37 ±40.23 | 30.65 ±6.05 | 0.26 ±0.01 | 0.20 ±0.2 | 0.29 ±0.04 | 0.22 ±0.03 | 118.24 ±9.72 | 167.33 ±14.86 |

***S11 Platinum group element (PGEs) biomining.*** Total concentration (in ng/mL, or ppb) extracted from the meteorite rock, percentage of element extraction compared to total concentration in the meteoritic material ($\%_{M}$), and percentage ratios with the non-biological controls ($\%_{NB}$), for each gravity condition (ISS = microgravity; Earth = Earth gravity) of ruthenium, palladium and platinum. NB = non-biological; SD = *S. desiccabilis*; PS = *P. simplicissimum*; CN = Consortium.

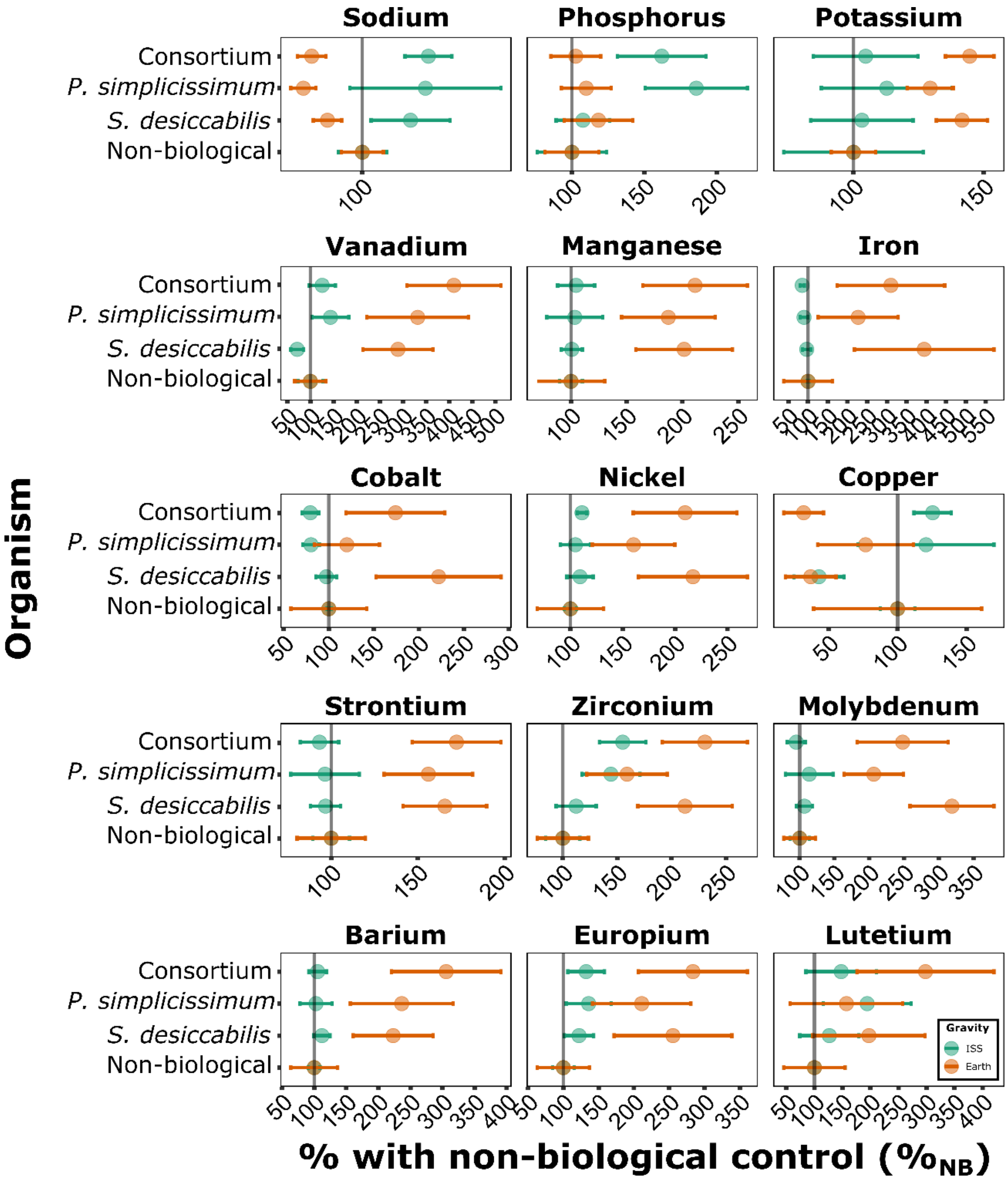


***S12*** Percentage ratios with the non-biological control ($\%_{NB}$) for further 15 elements whose bioextraction showed at least one p-value < 0.05 (ANOVA). Values are separated per gravity condition (ISS = microgravity, green circles, Earth = 1 x *g*, orange circles). Circles indicates mean values; error bars indicate standard error; grey vertical lines indicate 100% values (i.e., same bioleaching efficiency as non-biological samples).

| | | $\%_{NB}$ | | | | $\%_{M}$ | | | |
|---|---|---|---|---|---|---|---|---|---|
| | | **Non-biological** | ***S. desiccabilis*** | ***P. simpliciss.*** | **Consortium** | **Non-biological** | ***S. desiccabilis*** | ***P. simpliciss.*** | **Consortium** |
| Na | **ISS** | 100.00±6.41 | 112.83±10.51 | 116.79±20.09 | 117.57±6.24 | 0.57±0.07 | 0.64±0.09 | 0.66±0.14 | 0.67±0.08 |
| | **Earth** | 100.00±5.61 | 90.72±3.77 | 84.31±3.36 | 86.53±3.78 | 0.82±0.11 | 0.74±0.09 | 0.69±0.08 | 0.71±0.09 |
| P | **ISS** | 100.00±23.98 | 107.58±18.54 | 185.86±35.28 | 162.02±30.60 | 0.12±0.03 | 0.15±0.02 | 0.26±0.04 | 0.23±0.04 |
| | **Earth** | 100.00±18.59 | 118.30±23.72 | 109.84±17.17 | 102.63±17.22 | 0.18±0.03 | 0.21±0.04 | 0.19±0.03 | 0.18±0.03 |
| K | **ISS** | 100.00±26.79 | 103.22±19.65 | 112.77±25.14 | 104.67±20.06 | 1.13±0.23 | 1.16±0.09 | 1.27±0.18 | 1.18±0.10 |
| | **Earth** | 100.00±8.55 | 141.65±9.85 | 129.44±8.85 | 144.62±9.30 | 1.06±0.10 | 1.50±0.13 | 1.37±0.11 | 1.53±0.12 |
| V | **ISS** | 100.00±27.00 | 71.40±13.97 | 143.47±39.69 | 125.54±28.67 | NA | NA | NA | NA |
| | **Earth** | 100.00±33.98 | 289.49±75.64 | 332.18±109.81 | 410.51±101.8 | NA | NA | NA | NA |
| Mn | **ISS** | 100.00±10.21 | 100.76±9.45 | 103.37±25.16 | 104.55±16.58 | 0.0003±0.0000 | 0.0003±0.0000 | 0.0003±0.0001 | 0.0003±0.0001 |
| | **Earth** | 100.00±30.30 | 201.47±43.39 | 187.34±41.98 | 211.31±46.98 | 0.0002±0.0000 | 0.0003±0.0000 | 0.0003±0.0000 | 0.0003±0.0000 |
| Fe | **ISS** | 100.00±2.84 | 96.67±10.93 | 89.83±10.15 | 85.12±5.26 | 0.11±0.01 | 0.10±0.02 | 0.10±0.02 | 0.09±0.01 |
| | **Earth** | 100.00±61.61 | 394.35±176.82 | 227.03±101.55 | 309.94±136.26 | 0.02±0.01 | 0.10±0.02 | 0.06±0.01 | 0.08±0.01 |
| Co | **ISS** | 100.00±2.40 | 97.34±11.64 | 80.24±9.23 | 79.53±9.28 | 0.05±0.10 | 0.49±0.11 | 0.40±0.10 | 0.40±0.10 |
| | **Earth** | 100.00±42.01 | 221.93±69.40 | 120.03±36.20 | 174.04±54.83 | 0.21±0.10 | 0.46±0.10 | 0.25±0.05 | 0.36±0.10 |
| Ni | **ISS** | 100.00±5.89 | 109.18±12.58 | 104.75±14.25 | 111.02±4.68 | 0.15±0.01 | 0.16±0.02 | 0.16±0.02 | 0.17±0.01 |
| | **Earth** | 100.00±31.77 | 217.12±52.12 | 160.30±39.46 | 209.64±49.57 | 0.06±0.01 | 0.14±0.01 | 0.10±0.01 | 0.13±0.01 |
| Cu | **ISS** | 100.00±12.48 | 43.01±18.10 | 120.60±49.22 | 125.38±13.51 | 0.002±0.000 | 0.001±0.000 | 0.002±0.001 | 0.002±0.000 |
| | **Earth** | 100.00±60.88 | 37.07±18.33 | 76.83±34.74 | 31.92±14.42 | 0.014±0.006 | 0.005±0.001 | 0.010±0.002 | 0.004±0.001 |
| Sr | **ISS** | 100.00±10.55 | 96.73±8.75 | 96.41±19.79 | 93.22±11.09 | 0.39±0.06 | 0.37±0.05 | 0.37±0.09 | 0.36±0.06 |
| | **Earth** | 100.00±19.73 | 165.53±24.11 | 156.02±25.57 | 172.28±25.62 | 0.21±0.04 | 0.35±0.05 | 0.33±0.05 | 0.37±0.05 |
| Zr | **ISS** | 100.00±15.67 | 112.00±12.63 | 144.23±26.72 | 155.06±21.38 | 0.002±0.000 | 0.002±0.000 | 0.002±0.000 | 0.002±0.000 |
| | **Earth** | 100.00±23.54 | 212.33±43.33 | 159.03±37.02 | 230.62±39.13 | 0.001±0.000 | 0.002±0.000 | 0.001±0.000 | 0.002±0.000 |
| Mo | **ISS** | 100.00±13.90 | 106.77±11.63 | 113.83±34.27 | 95.05±12.86 | 0.03±0.01 | 0.03±0.01 | 0.03±0.02 | 0.03±0.01 |
| | **Earth** | 100.00±22.93 | 319.16±60.41 | 206.61±42.90 | 248.24±65.63 | 0.01±0.01 | 0.03±0.01 | 0.02±0.01 | 0.02±0.01 |
| Ba | **ISS** | 100.00±9.13 | 112.00±12.63 | 102.73±24.92 | 105.31±13.78 | 0.11±0.02 | 0.13±0.02 | 0.12±0.03 | 0.12±0.02 |

| | | | | | | | | | |
|---|---|---|---|---|---|---|---|---|---|
| | **Earth** | 100.00±36.61 | 223.05±62.37 | 236.39±80.09 | 305.66±85.39 | 0.05±0.02 | 0.12±0.02 | 0.13±0.03 | 0.16±0.03 |
| Eu | **ISS** | 100.00±15.20 | 122.22±20.78 | 135.81±31.78 | 132.31±26.00 | 0.003±0.000 | 0.004±0.001 | 0.004±0.001 | 0.004±0.001 |
| | **Earth** | 100.00±36.97 | 255.23±83.41 | 210.97±69.65 | 283.56±77.18 | 0.001±0.000 | 0.003±0.001 | 0.003±0.001 | 0.004±0.000 |
| Lu | **ISS** | 100.00±54.73 | 126.63±52.78 | 194.13±78.51 | 147.77±63.01 | 0.004±0.001 | 0.004±0.001 | 0.006±0.001 | 0.006±0.001 |
| | **Earth** | 100.00±54.70 | 197.23±100.03 | 157.16±100.53 | 294.13±122.21 | 0.002±0.001 | 0.002±0.001 | 0.003±0.001 | 0.003±0.001 |

***S13*** Percentage ratios with the non-biological controls ($\%_{NB}$), for each gravity condition (ISS = microgravity; Earth = Earth gravity) of the further 15 elements whose at least one biologically relevant pairwise comparison provided a p-value <0.05 (mean±standard error). Elements are listed in order of atomic number.

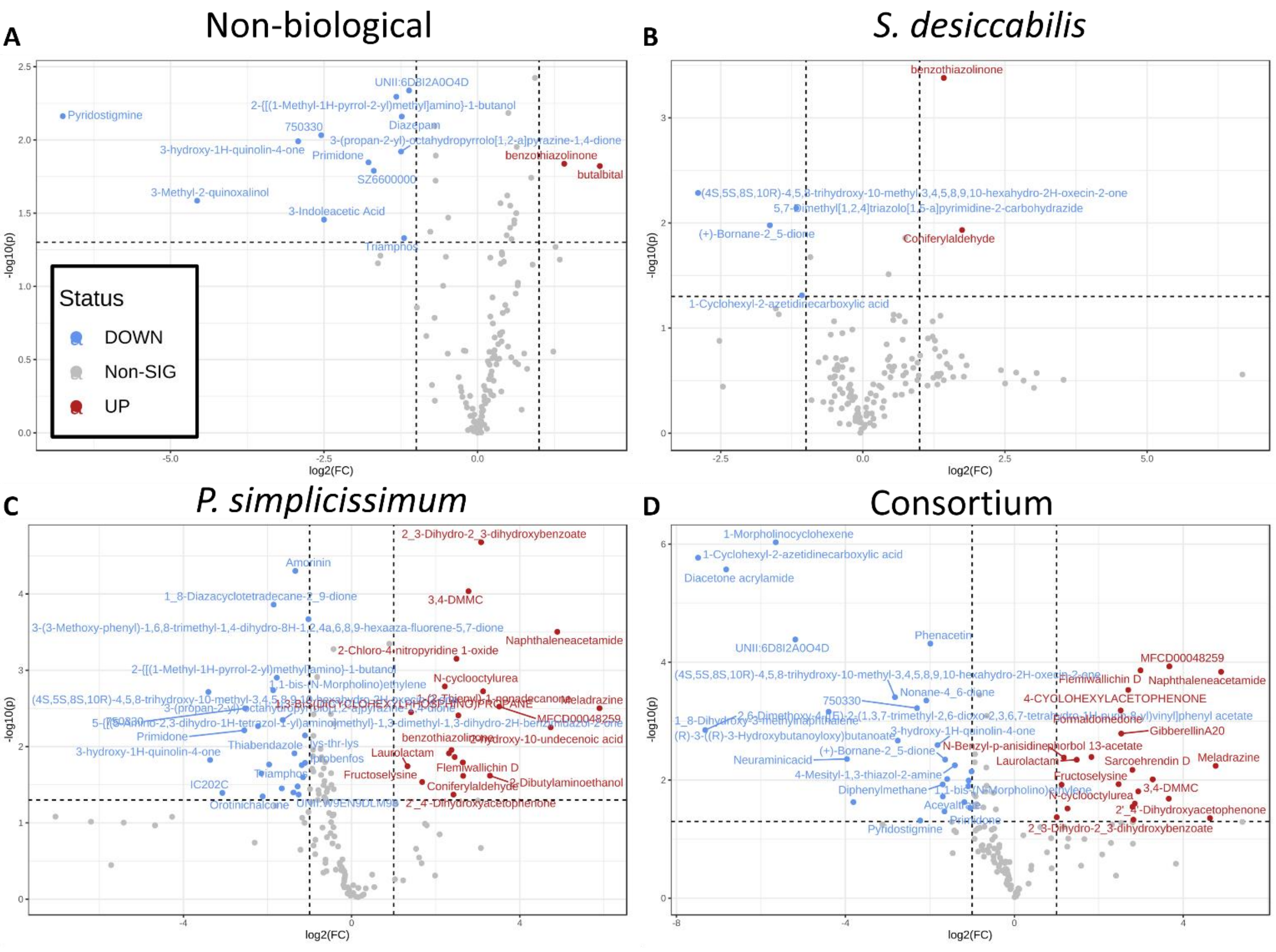


***S14*** Volcano plots showing up- or downregulations of features in space compared to Earth. Comparisons are shown for A) non-biological, B) *S. desiccabilis*; C) *P. simplicissimum* and D) Consortium samples.

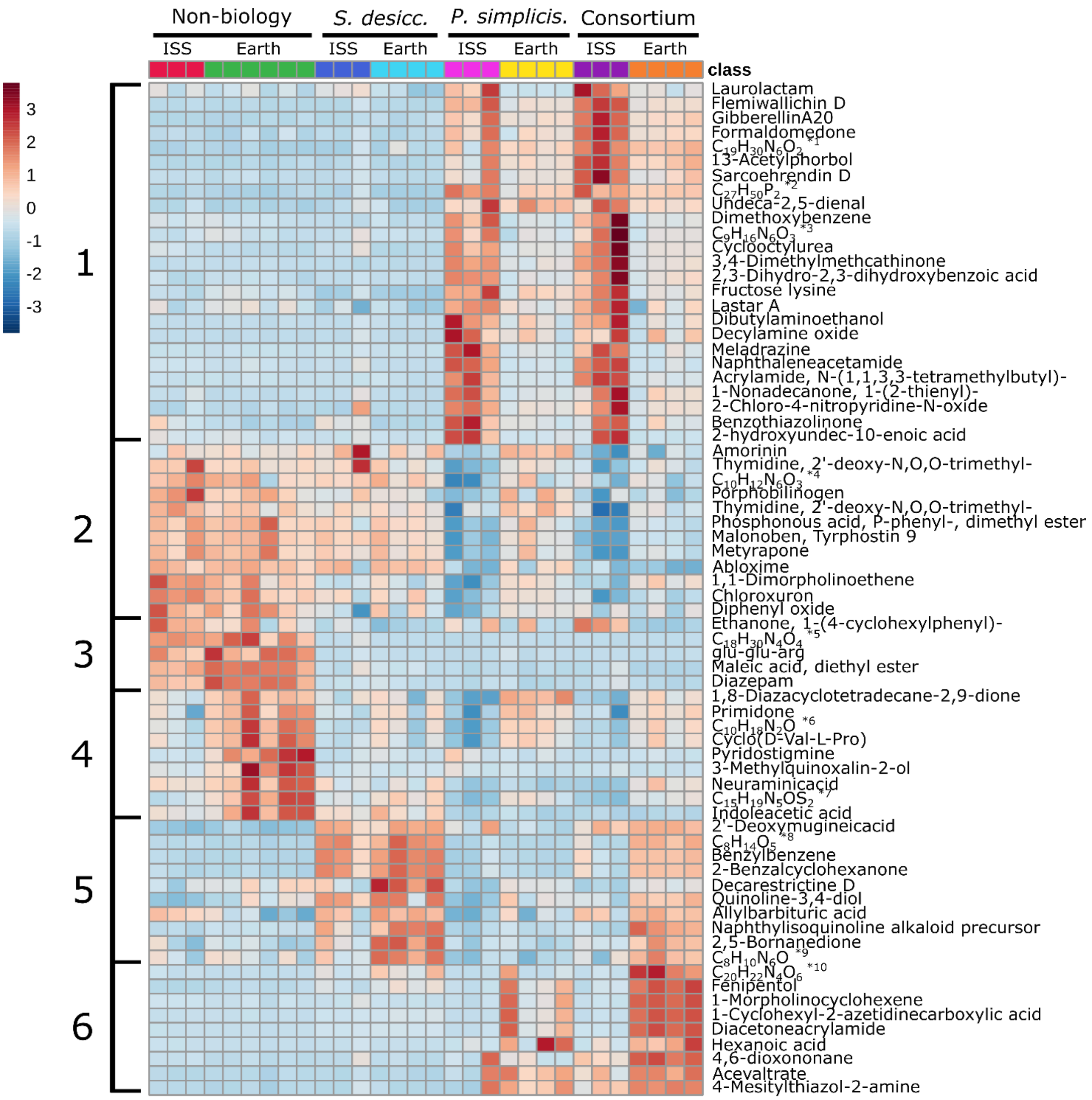


*[1] N-[2-(1-Cyclohexen-1-yl)ethyl]-4,6-di(4-morpholinyl)-1,3,5-triazin-2-amine
*[2] Phosphine, 1,3-propanediylbis[dicyclohexyl-
*[3] 2-{4-[(Z)-(4-Amino-1,2,5-oxadiazol-3-yl)(hydroxyimino)methyl]-1-piperazinyl}ethanol
*[4] 7-Amino-1,3-dimethyl-2,4-dioxo-1,2,3,4-tetrahydropyrido[2,3-d]pyrimidine-6-carbohydrazide
*[5] 2,3-Bis(4-morpholinylmethyl)-4a,5,6,7,8,8a-hexahydroquinoxaline 1,4-dioxide
*[6] 1-Butanol, 2-[[(1-methyl-1H-pyrrol-2-yl)methyl]amino]-
*[7] Acetamide, 2-[(8,9-dimethylthieno[3,2-e]-1,2,4-triazolo[4,3-c]pyrimidin-3-yl)thio]-N,N-diethyl-
*[8] (R)-3-((R)-3-Hydroxybutanoyloxy)butanoate
*[9] 5,7-Dimethyl[1,2,4]triazolo[1,5-a]pyrimidine-2-carbohydrazide
*[10] 2,6-Dimethoxy-4-[(E)-2-(1,3,7-trimethyl-2,6-dioxo-2,3,6,7-tetrahydro-1H-purin-8-yl)vinyl]phenyl acetate

***S15*** Heatmap showing the top 70 features measured in the metabolomics analysis, clustered per gravity condition and organism. Each column represents a single sample. Numbers on the left represent 6 visual clusters suggesting metabolomic patterns dependent on the organism and/or the gravity condition. Cluster 1 represents features expressed almost solely in the presence of *P. simplicissimum* in space. Cluster 2 shows features present in all samples except those containing the fungus (alone or in Consortium) on the ISS, potentially indicating features consumed by the fungi when in microgravity conditions. Cluster 3 shows features present in the non-biological control samples regardless of gravity condition, while cluster 4 shows features mostly present in the non-biological control samples in Earth gravity. Cluster 5 shows features highly associated with the presence of *S. desiccabilis*. Complementary to Cluster 1, Cluster 6 represents features solely produced in the presence of the fungus in Earth gravity. Interestingly, the observation of these clear patterns, together with the separate clusters observed in Figure 6D, suggests that the contaminant species (S3) had no or minor effects on the metabolomics results. Features with long names were indicated with their chemical formula, with asterisks showing the complete names at the bottom of the figure.

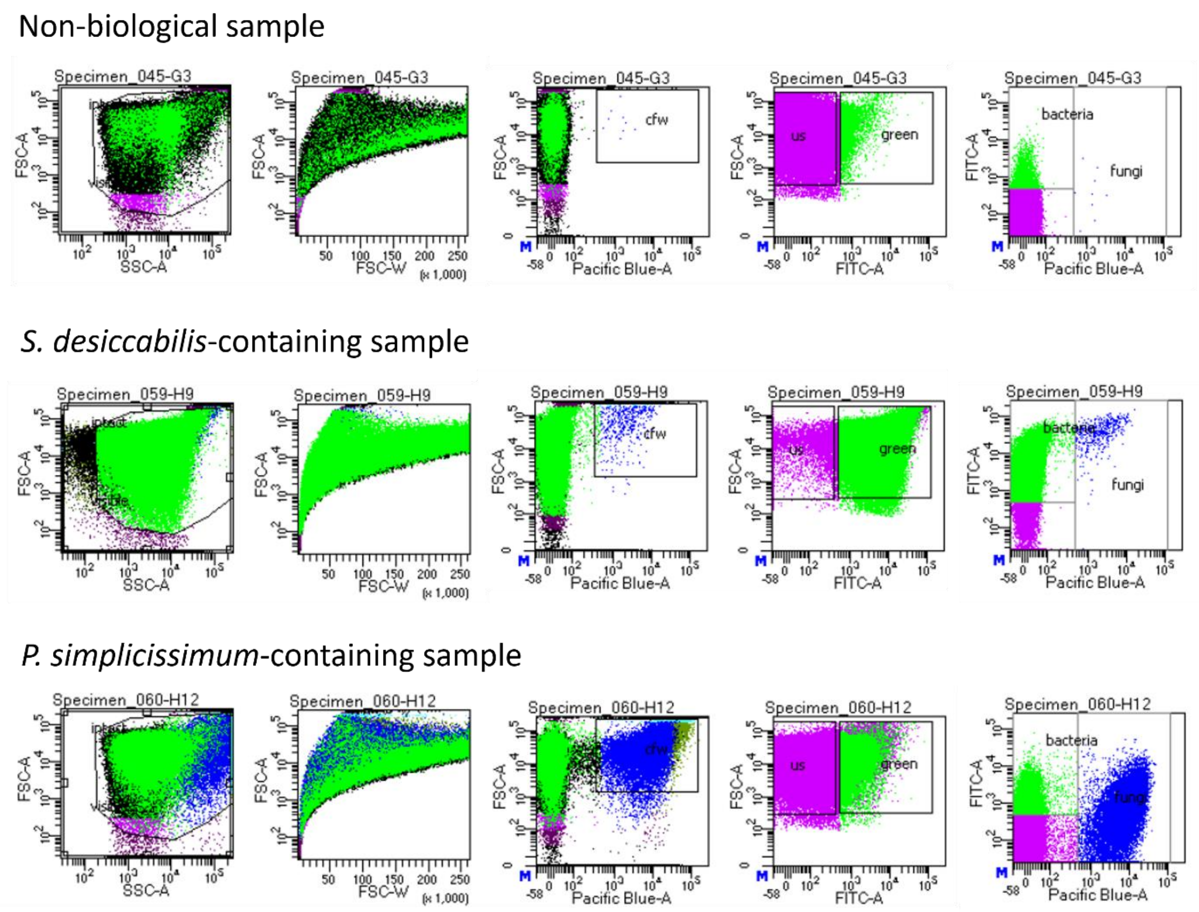


***S16*** Gating strategy for the flow cytometry analysis, constructed using the software BD FacsDiva 8.0.1, to distinguish bacterial from fungal cells. Three representative samples are shown here (top: a non-biological space control; middle: a *S. desiccabilis* containing sample; bottom: a *P. simplicissimum* containing sample. The events reported in this manuscript were obtained from the *fungi* and *bacteria* gating. The Pacific Blue-A filter was used to visualise calcofluor white (*cfw*, fungi-specific dye), while FITC-A filter was used for backlight green (*green*, bacteria-specific stain). *us* gating indicates unstained particles.

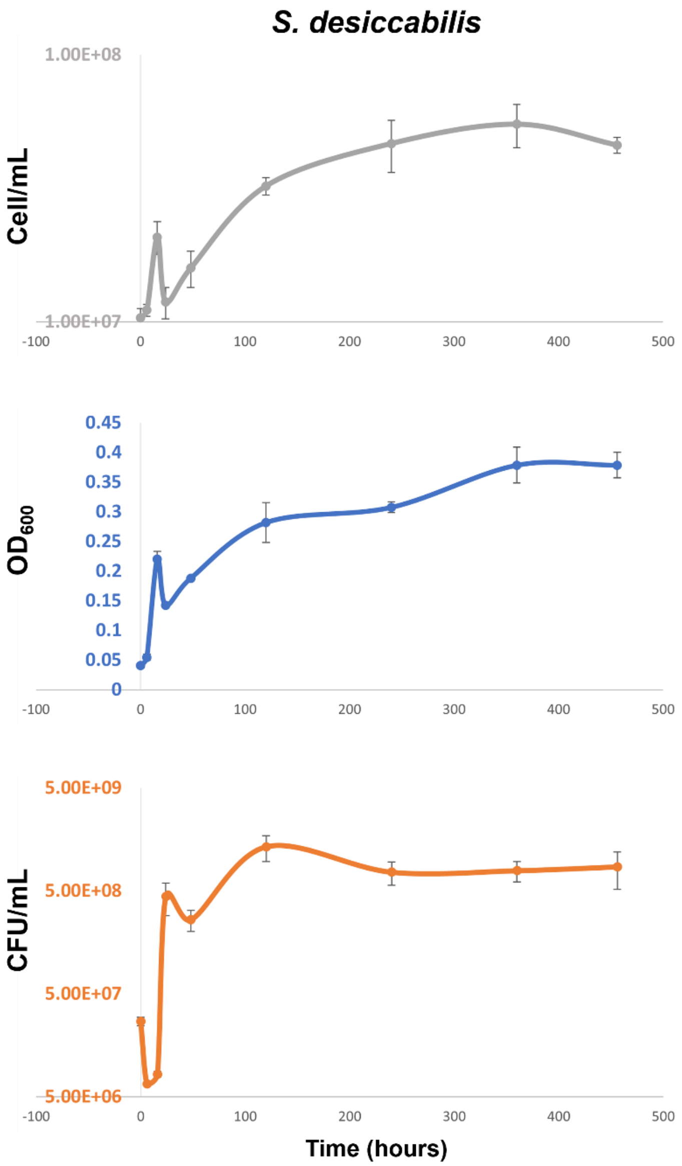


***S17*** Growth curve for *S. desiccabilis* showing comparison between the three methods for final cell concentration used in this work. *S. desiccabilis* growth curve was determined in 19 days (456 hours), which is the timeline used for the BioAsteroid experiment. Flow cytometry (cell/mL, in grey), $OD_{600}$ (in blue) and CFU/mL (in orange), are shown. Points represent average values, error bars represent standard errors.

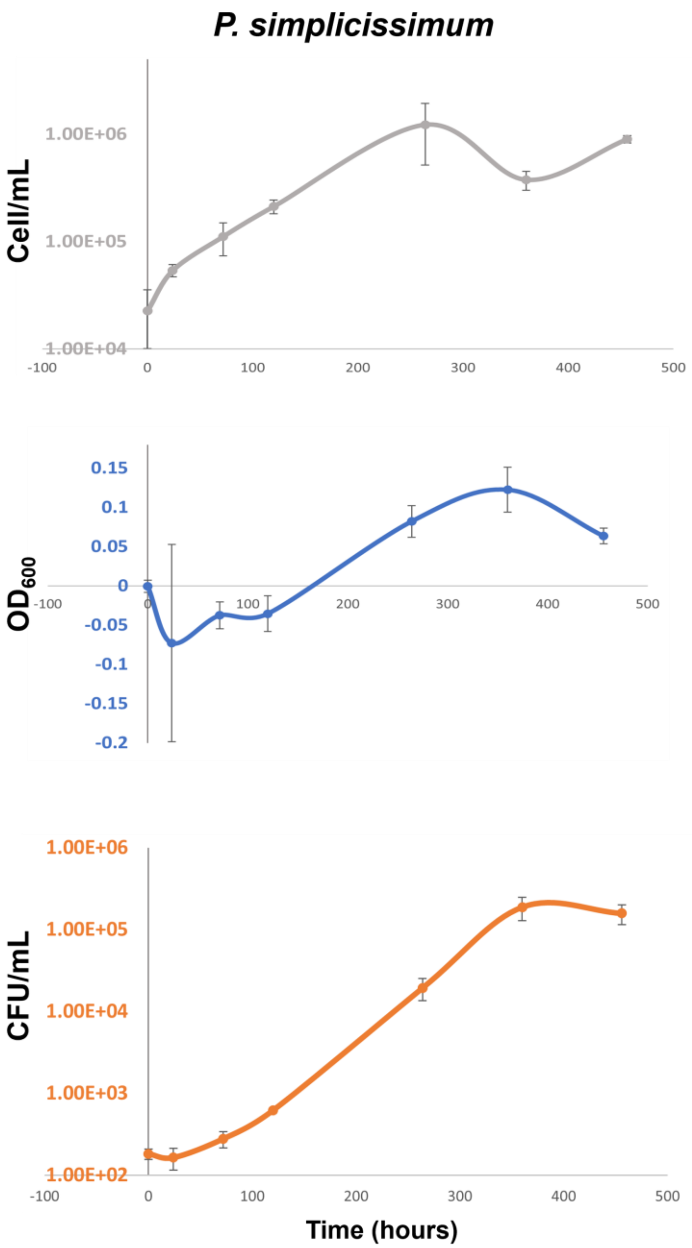


***S18*** Growth curve for *P. simplicissimum* showing comparison between the three methods for final cell concentration used in this work. *P. simplicissimum* growth curve was determined in 19 days (456 hours), which is the timeline used for the BioAsteroid experiment. Flow cytometry (cell/mL, in grey), $OD_{600}$ (in blue) and CFU/mL (in orange), are shown. Points represent average values, error bars represent standard errors.

### References for supplementary information


Cockell, C. S. et al. Microbially-Enhanced Vanadium Mining and Bioremediation Under Micro- and Mars Gravity on the International Space Station. *Front. Microbiol.* **12**, 663 (2021).

Cockell, C. S. et al. Space station biomining experiment demonstrates rare earth element extraction in microgravity and Mars gravity. *Nat. Commun.* **11**, 1–12 (2020).

Petrikkou E, Rodríguez-Tudela JL, Cuenca-Estrella M, Gómez A, Molleja A, Mellado E. Inoculum standardization for antifungal susceptibility testing of filamentous fungi pathogenic for humans. *J Clin Microbiol.* 2001 Apr;**39**(4):1345-7. doi: 10.1128/JCM.39.4.1345-1347.2001.
